# The activation-deactivation cycle of human PIEZO1

**DOI:** 10.64898/2026.11.12.766215

**Authors:** Joana Massa, Benedikt Frieg, Christian Tyrchan, Oliver Koch

## Abstract

PIEZO1 is a cation channel that is primarily activated through mechanical force, and activation has been shown to be influenced by membrane curvature. PIEZO1 is ubiquitously expressed and plays critical roles in various patho-/physiological processes such as inflammatory diseases or red blood cell volume regulation. Here we combine coarse-grained and atomistic simulations in which human PIEZO1 is embedded in a curved membrane and subjected to simulations under controlled membrane tension and a physiological transmembrane potential, allowing the gating cycle to be reconstructed at atomic resolution. Systematic pressure adaptation identifies an activation regime in which the channel flattens progressively, and ion permeation quantitatively reproduces the experimental conductance of 36 pS. Furthermore, the R2456-E2117 salt bridge couples pore lipid retention to the anchor domain and disrupting it reproduces the phenotypes of the disease-associated R2456H and R2456C variants. PIEZO activation is transmitted through an interconnected network of contact rearrangements that ultimately reshapes the yoda1 binding pocket at the THU8/THU9 interface. This state-dependent remodelling rationalises the pharmacology of yoda1 and offers a framework for the structure-based design of PIEZO1 modulators.

## Introduction

Together with PIEZO2, PIEZO1 forms a unique family of ion channels inherently sensitive to pressure changes, which is essential for, e.g. translating the sensation of touch in mammals. PIEZO1 is found in nearly all tissues and plays a role in processes involving mechanical force, such as neuronal development, cell size regulation, and immune system activity. Dysregulation of PIEZO1 can lead to various diseases. In humans, loss-of-function (LOF) mutations in PIEZO1 cause Lymphatic Malformation 6, while gain-of-function (GOF) mutations result in dehydrated hereditary stomatocytosis 1. Notably, the E756del mutation increases malaria resistance but leads to iron overload in red blood cells (RBCs). There, PIEZO1 activation causes calcium influx, which triggers activation of KCa3.1 and leads to cell shrinkage.^1^

PIEZO1 is a homotrimeric protein that is primarily activated by mechanical force. The channel consists of nine transmembrane helical units (THUs) consisting of four transmembrane helices (TMs) each, totalling 36 helices for the blade domain (Fig. 1). Additionally, the ion conducting pore consists of two helices called inner and outer helices (IH and OH). The nine THUs make up the blades that surround the central pore. In experimental structures of mammalian PIEZO1, the first three THUs are not resolved as they are believed to be too flexible.^2–4^ Previous models of channel activation postulate that the Beam, Clasp and extracellular loop (ECL) domains are responsible for translating the pressure from the ends of the blade towards the centre of the channel. Above the centre, there is a Cap domain that influences ion flow through the pore.^1,3,5^

**Fig. 1:**
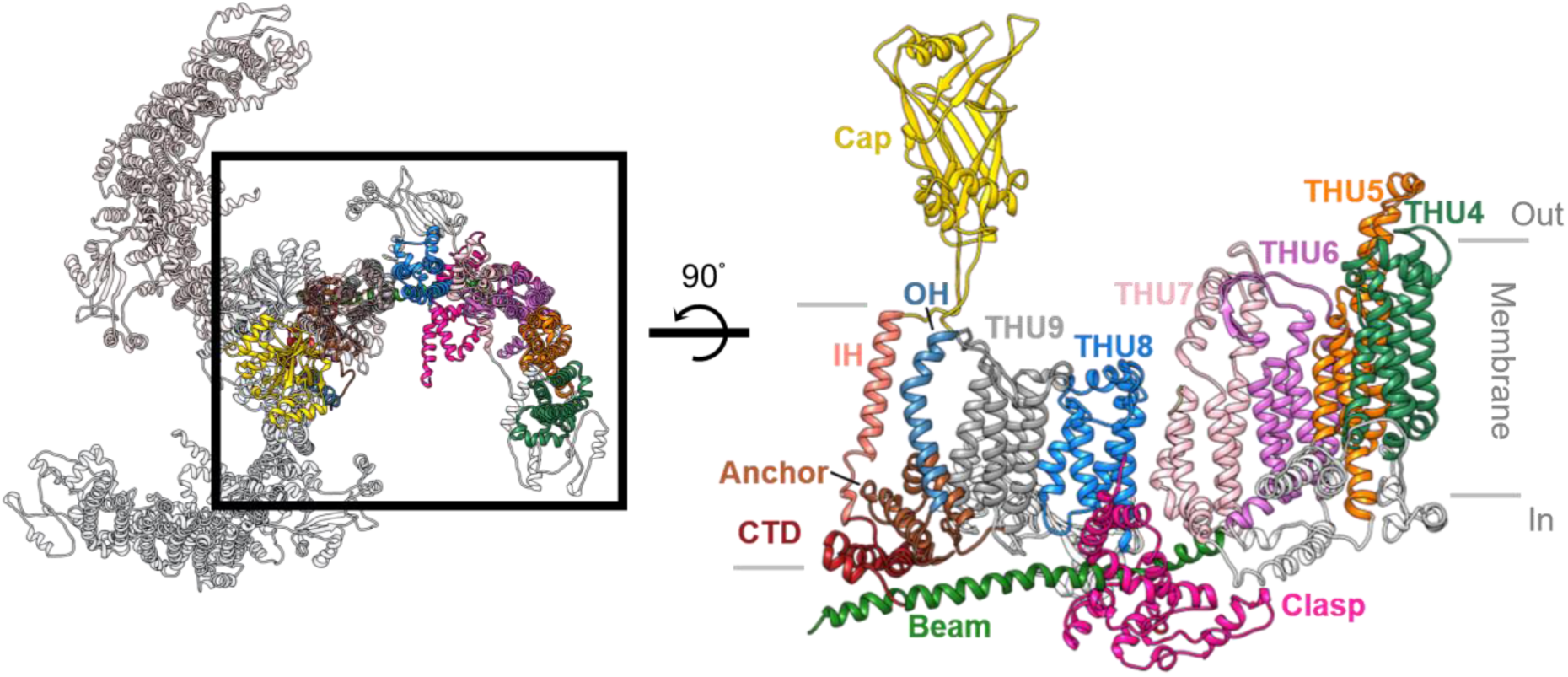
Top view of PIEZO1 (left). On the right a detailed structural overview of one human PIEZO1 subunit as modelled in this study is shown.

To enable simulation of large-scale biomolecular processes as opposed to all-atom (AA) models, compromises have to be made for PIEZO1 by replacing atomistic detail with lower-resolution, coarse-grained (CG) beads.^6–8^ By reducing the number of degrees of freedom, CG models decrease the computational cost while also reducing entropic contributions. In addition, the simplified interaction potentials smooth the underlying energy landscape, allowing simulations to escape local energy minima that can hinder efficient sampling in AA simulations.^9^ Consequently, CG approaches are well suited for studying large systems and long-timescale phenomena, although this increased efficiency comes at the expense of atomic-level detail.

One of the most widely used CG force fields is MARTINI. It employs a one-to-four mapping scheme, in which, on average, four heavy atoms together with their associated hydrogen atoms are represented by a single coarse-grained bead. Combined with its modular building-block approach for describing chemical groups, this mapping facilitates straightforward conversion between coarse-grained and all-atom representations.^10^

Here, we aim to combine the speed-up and robustness of CG simulations with the atomistic details provided by all-atom representations (Fig. 2). Previous computational studies investigated ion conductance and provided important insights into PIEZO1’s mechanosensitivity, revealing roles for membrane tension, blade flattening and pore dilation^3,11^ as well as interblade interactions^12^, lipid rearrangement^13^, and linker mechanics in channel gating^5^. Studies of ligand binding further demonstrated that the small molecule yoda1 binds to the channel at the THU8/9 interface modulating ion conductance and the channel’s mechanosensitivity.^14–16^ However, most previous simulations were based on mouse PIEZO1.^13^ Therefore, the membrane stability and large-scale mechanical coupling of human PIEZO1 remain poorly studied *in silico*. The recent elucidation of a human PIEZO1 structure provides an opportunity to address these limitations and comprehensively link membrane curvature with changes on the atomic level. Consequently, we use molecular dynamics (MD) simulations of human PIEZO1 to investigate its structural and membrane stability and to characterise structural changes that may underlie mechanosensitive gating and ligand binding.

**Fig. 2:**
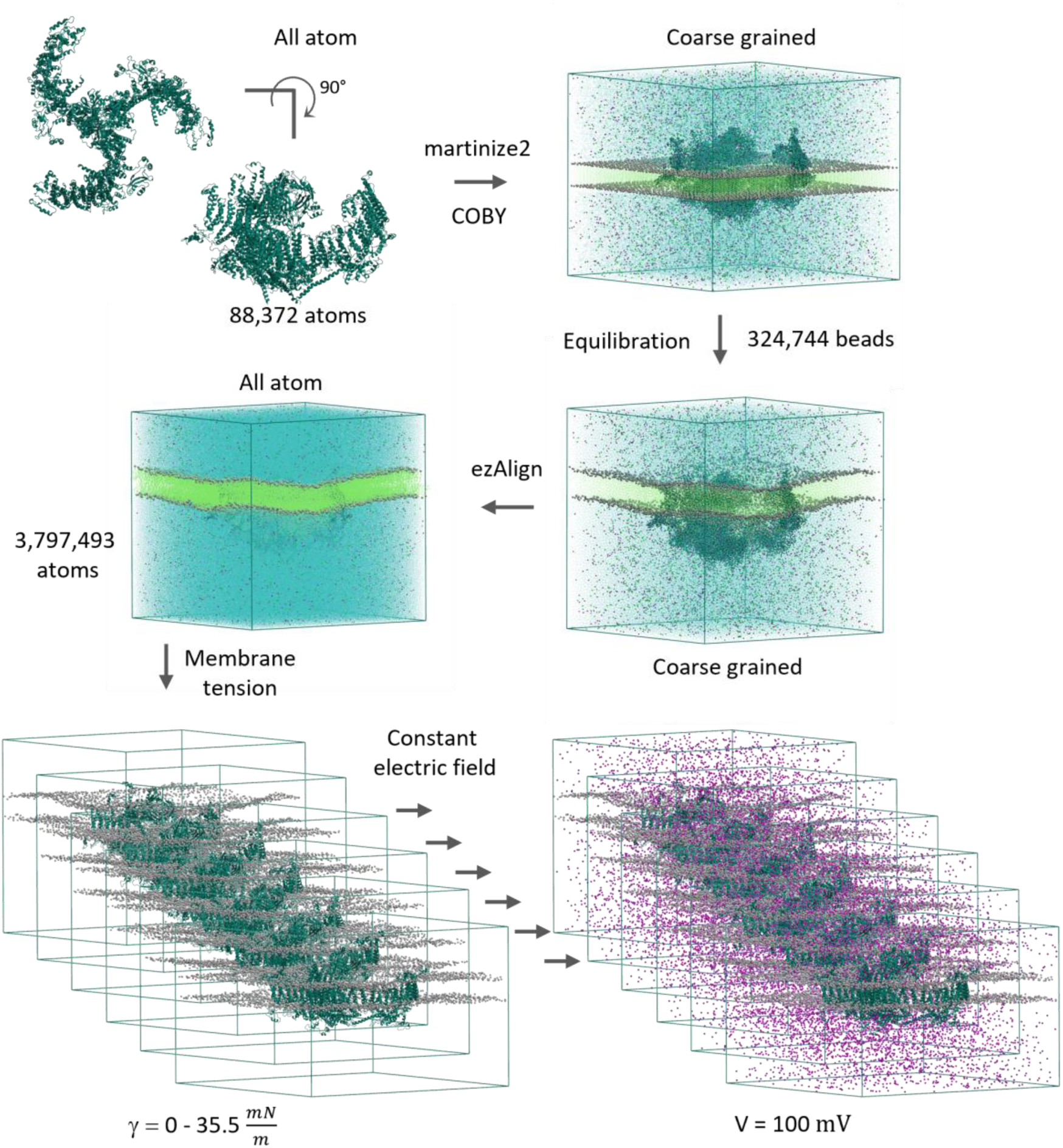
The human PIEZO1 channel consists of three subunits that from the top each show a characteristic blade-like structure. From the side, the curvature is visible. In this study, we simulated the activation pathway starting from the curved, closed structure towards an open, flattened channel. The workflow employed in this work is illustrated in the following: Starting out from the all-atom protein model, it was converted to coarse-grained with martinize2. Then, the coarse-grained protein model was inserted into a flat membrane and water and ions were added using the toolkit COBY. After equilibration, the system represents a coarse-grained curved, inactive channel in a buckle-like membrane. This system was converted back to all atom using ezAlign. Then the activation mechanism was investigated by application of i) different pressures, i.e. membrane tensions (γ) and ii) a constant electric field (V).

## Methods

### Preparation of human PIEZO1 model

The protein was modelled based on the human PIEZO1 structure with the PDB ID 8YEZ.^2^ First, all missing loops were built by using the “Homology Builder” tool of the Molecular Operating Environment (MOE) v2024.0602 using default settings.^17^ Then, the model with the best GB/IV score was selected for further refinement. The protonation was adjusted to a pH of 7.4 with the “Protonate 3D” tool and then the model was energy minimized using the AMBER14:EHT^18^ force field while tethering the atoms. A deviation of 1.5 Å was allowed. In the resulting model all loops except the intracellular loop connecting the Beam and Clasp region were included. Specifically, residues 1,371 to 1,512 were deleted as this loop was modelled as a long and loopy strand crossing through the pore. However, these residues would have formed the Central and Lateral Plug and Latch regions, which were postulated to influence ion flow through the intracellular fenestration. Therefore, our model more closely resembles the PIEZO1.1 isoform, that lacks both Plug domains and shows an increased single-channel conductance and mechanosensitivity compared to wild-type (WT).^19^ Structurally the three subunits of our model consist of two protein chains each. Similar to the experimental template, the THU1-3 were not modelled. Therefore, residues 1-569 are missing. In total, one subunit consists of 1,810 residues; the whole protein model has 5,430 residues and 88,372 atoms. In case of the mutated channel systems, R2456 was changed to cysteine or histidine using the MOE “Protein Builder” tool. The histidine was protonated at the ε-nitrogen. The R2456C and R2456H systems include 88,342 and 88,351 protein atoms, respectively.

### Coarse-grained modelling of curved PIEZO1

The prepared all-atom structures of WT and mutated PIEZO1 was converted to their coarse-grained (CG) Martini3^20,21^ representation using martinize2^22^. All following workflow steps were followed for all three systems separately. For modelling of the protein secondary and tertiary structure, an elastic network model of 7 Å and 9 Å for the lower and upper cutoff distance was used, respectively. This curved CG protein model was inserted into a flat CG POPC membrane using the tool COBY^23^. A rectangular box with a size of 350 x 350 x 300 Å^3^ was used. For water and salt the default molarity values of 55.56 mol L^-1^ and 0.15 mol L^-1^ were used which resulted in a system with a total of 324,744 beads. In detail, the WT system includes 1,802 POPC (upper leafleat), 1,819 POPC (lower leaflet), 262,481 water, 2,847 sodium and 2,847 chloride beads. All simulations were conducted using GROMACS v2024.2^24^. First, the system was energy minimised for 5000 steps using the steepest descent algorithm. Afterwards, a second energy minimisation was performed for 5000 steps using the conjugate gradient algorithm. For all simulation steps, Coulomb and van-der-Waals (vdW) cutoffs of 11 Å were used and periodic boundary conditions in all directions were enabled. The Coulomb-type reaction field with a relative dielectric constant (epsilon-r) of 15 was used. To adapt the initial flat membrane to the curved protein shape, the protein backbone beads were restrained during the CG simulation. After energy minimisation, the system was simulated for 1 µs with a time step of 20 fs. In this simulation, the temperature and pressure were set to 300 K and 1 bar separately controlled for the bilayer (lipids and protein) and solvent with the V-rescale thermostat (τt = 1 ps) and Berendsen barostat (τp = 3 ps), respectively. Semi-isotropic pressure coupling and a compressibility of 4.5e^-5^ was used. The CG simulations were run on a machine with a NVIDIA GeForce 4090 GPU and a 24-core AMD Ryzen Threadripper Pro 7965wx CPU reaching a performance of 931 ns day^-1^. The last frame of this simulation was then converted to all-atom using the tool ezAlign^25^, which resulted in system with 3,797,493 atoms for the WT protein. Details about the all-atom composition of the three systems (WT, R2456C and R2456H) are found in the Supporting Information Table S1.

### All-atom simulations with different pressures

The all-atom models, resulting from the back mapping procedure with ezAlign^25^, were further energy minimised and then equilibrated using six short simulations while gradually releasing restraints on the protein and membrane. The procedure is similar to the default procedure from CHARMM-GUI^26^. For the protein and lipids, the CHARMM36^27^ force field was used. The parameters of water were taken from the TIP3P^28^ water model. In all all-atom simulations, the Coulomb and vdW cutoffs were set to 1.2 Å. VdW switching was enabled at 1.0 Å. Coulomb interactions were calculated using Particle Mesh Ewald (PME)^29^. Hydrogen bonds were constrained using the LINCS^30^ algorithm.

First, the system was energy minimised for 5000 steps using the steepest descent algorithm, while restraining protein backbone atoms (4000 kJ mol^-1^ nm^-2^). In the first equilibration simulations this restraint was kept, and the time step was 1 fs. In this and following simulations, the temperature was maintained at 300 K with the V-rescale thermostat (τt = 1 ps). This was done separately for the three groups: protein, membrane and solvent. The translational velocity was removed every 100 steps for the two groups protein_membrane and solvent. After the first simulation of 125 ps, the protein restraints were lowered to 2000 kJ mol^-1^ nm^-2^ and further simulated for 125 ps. In the following simulations, semi-isotropic pressure coupling was enabled. For this, the C-rescale barostat (τp = 5 ps, compressibility 4.5e^-5^) and a reference pressure of 1 bar was used. In the third 125 ps simulation, the 1 fs time step was still used and the restraints were lowered to 1000 kJ mol^-1^ nm^-2^. All following simulations use a time step of 2 fs. In the fourth, fifth and sixth simulation the protein backbone restraints were lowered from 500 to 200 and 50 kJ mol^-1^ nm^-2^ and each time the system was simulated for 500 ps.

The last frame of the sixth equilibration simulation was then used as the starting point for replicate production simulations at different pressures. For efficient simulation, all following production simulations were run on CPU only nodes of the Fritz HPC cluster (NHR@FAU). The best performance was achieved by using seven nodes (each with two 36 core Intel Xeon Platinum 8360Y CPUs) with in total 504 cores and allocating 88 threads for PME calculation balancing resource usage and computation speed. Dynamic load balancing was enabled. This resulted in an average performance of ∼10 ns day^-1^. In general, five replicas per pressure and system were simulated for 50 ns and coordinates were saved every 100 ps.

The semi-isotropic barostat was set to Pxy 1, -1, -3, -5, -8, -10, -20, -30 and -40 bar and Pz 1 bar. This ensemble pressure can be used to estimate the applied membrane tension γ with the following equation:

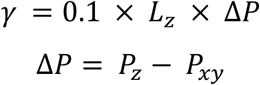

The box z-dimension (Lz) of the system after equilibration was used to estimate the membrane tensions (Table 1).

**Table 1:** Estimated membrane tension.

| **System No.** | **P<sub>z</sub> [bar]** | **P<sub>xy</sub> [bar]** | **L<sub>z</sub> [nm]** | **γ [mN m<sup>-1</sup>]** | **Acronym** | **Replicas** |
|---|---|---|---|---|---|---|
| **1** | 1 | 1 | 32.3 | 0 | P_1 | 5 |
| **2** | 1 | -1 | 32.3 | 6.5 | P_m_1 | 5 |
| **3** | 1 | -3 | 32.3 | 12.9 | P_m_3 | 5 |
| **4** | 1 | -5 | 32.3 | 19.4 | P_m_5 | 5 |
| **5** | 1 | -8 | 32.3 | 29.1 | P_m_8 | 5 |
| **6** | 1 | -10 | 32.3 | 35.5 | P_m_10 | 5 |
| **7** | 1 | -20 | 32.3 | 67.8 | P_m_20 | 5 |
| **8** | 1 | -30 | 32.3 | 100.1 | P_m_30 | 1 |
| **9** | 1 | -40 | 32.3 | 132.4 | P_m_40 | 1 |
| **10** | 1 | -5 | 32.4 | 19.4 | R2456C | 5 |
| **11** | 1 | -5 | 32.5 | 19.5 | R2456H | 5 |

### All-atom constant electric field simulations

Five replicas of the 1, -1, -3, -5, -8 and -10 bar systems were selected for further simulation. In this next set of simulations an NVT ensemble was used to keep the box volume constant to enable the application of a constant electric field. Therefore, only the V-rescale thermostat was used and no pressure coupling was enabled. Otherwise, the simulation parameters as described for the varied pressure production simulations were kept. As the electric field depends solely on Lz^26^, a potential corresponding to 100 mV was applied throughout the whole simulation using the mdp file setting “electric-field” from GROMACS (Table 2).

**Table 2:** Overview over successfully simulated systems with an applied constant electric field, i.e. without ruptured membranes.

| **System No.** | **Sim. Time**<br>t **[ns]** | V **[mV]** | L<sub>z</sub> **[nm]** | **Acronym** | **Replicas** |
|---|---|---|---|---|---|
| **1** | 210 | 100 | ~31 | P_1_cef | 5 |
| **2** | 210 | 100 | ~30 | P_m_1_cef | 5 |
| **3** | 210 | 100 | ~29 | P_m_3_cef | 5 |
| **4** | 210 | 100 | ~27 | P_m_5_cef | 5 |
| **5** | 210 | 100 | ~26 | P_m_8_cef | 3 |
| **6** | 210 | 100 | ~25 | P_m_10_cef | 1 |
| **7** | 210 | 100 | ~27 | R2456C_cef | 5 |
| **8** | 210 | 100 | ~27 | R2456H_cef | 5 |

The ion conductance (G) was calculated by counting the number of Na^+^ permeation events (N) in the simulation time (t). The first 10 ns of each simulation were disregarded for this calculation. Permeation events were defined as the crossing of a boundary inside the ion pore in the centre of PIEZO1. The boundary defined in the box z dimension was set to the centre of mass (COM) of the backbone atoms of the pore residues L2435 to S2460. Only unique crossing events under the transmembrane potential (V) were taken into account for the calculation, as follows:

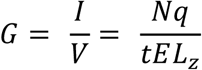

where q is the charge of an ion, E is the effective electric field and Lz is the box size in the z direction. The density map of ions, lipids and water was calculated using MDAnalysis.^31^ Individual box sizes and conductances of the replicas are found in the Supporting Information Table S2 and S3.

### Deactivation simulations

Selected simulations were performed using a third simulation approach (Table 3). Semi-isotropic pressure coupling was enabled again, resulting in an NPT ensemble. For this the C-rescale barostat with a τp of 5 ps and a compressibility of 4.5e^-5^ was used. Other simulation parameters were identical to those described before.

**Table 3:** Overview of deactivation simulations.

| **System No.** | **Sim. Time**<br>**t [ns]** | **P<sub>z</sub> [bar]** | **P<sub>xy</sub> [bar]** | **Acronym** | **Replicas** |
|---|---|---|---|---|---|
| **1** | 100 | 1 | 1 | P_5_1bar | 5 |
| **2** | 50 | 1 | 1 | P_m_10_bar | 1 |
| **4** | 100 | 1 | 1 | R2456H_1bar | 5 |

### Simulation analysis

The simulation trajectories were analysed, and plots were generated using in-house Python scripts relying on pre-existing libraries, mainly seaborn, Matplotlib, numpy, pandas, MDAnalysis and ProLIF.^31,32^ Area-per-lipid, area and membrane thickness were calculated using FATSLiM.^33^ The curvature analysis was done by custom scripts that were based on the MemCurv package.^34,35^ Lipid-protein interactions were analysed using ProLint v2.^36^ MD simulation systems and proteins were visualised using UCSF Chimera and VMD2.^37,38^ For contact analysis, Contact Map Explorer v0.7.0 was used which builds on MDTraj.^39,40^ Analysis of the pocket shape and properties of the THU8/9 interface was done using fpocket.^41^ The pocket shapes in Fig. 9 were generated by running fpocket on the yoda1 docked poses using the -a flag to identify the ligand chain.

### Protein-Ligand Docking

For the selection of the representative frame for docking the trajectories of P_1_cef, P_m_5_cef and P_m_10_cef were clustered on the backbone atoms of the THU8/9 interface (residues 1234-5678) of a single subunit using MDANCE and its k-means NANI implementation.^42^ For the concatenated trajectories this resulted in four to five clusters and the most representative frame of each cluster was used as input for ensemble docking with GOLD.^43^ All extracted structures were aligned on the backbone atoms of the largest cluster’s representative structure prior to docking. The structure of yoda1 was constructed using MOE and its builder utility. For further preparation, the conformation was energy minimised using the MMFF94x.^44^ The binding site was determined as a sphere with a 10 Å radius approximately in the middle of previously identified residues involved in binding of yoda1 (V1714, F1715, A1718, R1724, A2075, A2078 and R2082).^15,16,45^ Using an automatic search efficiency of 200%, 100 poses were generated, which were scored with ChemPLP.^46^ The resulting binding mode was visually inspected in UCSF Chimera.

### Ligand strain energy calculation

Ligand strain analysis complements docking scores by quantifying the internal energetic cost a ligand pays to adopt its bound conformation. Ligand strain energies of the docking poses were computed with the Strain Energy Calculation and Rescoring module in Schrödinger MacroModel (release 2025-03). For each ligand conformation, a tightly restrained minimization and an unrestrained minimization were performed, and the resulting energy difference was used to define the strain penalty. Minimizations were carried out in implicit water solvation. Cartesian restraints were applied to the bound-state minimization, with a half-width of 0.3 Å and a force constant of 120 kcal mol⁻¹ Å⁻². A constant energy offset of 4.00 kcal mol⁻¹ was subtracted from the raw strain, and the resulting value was scaled by a penalty factor of 0.25 before being added to the docking score. No conformational search for global minima was performed.

## Results & Discussion

### Coarse-grained-to-atomistic modelling captures curvature-induced membrane remodelling

Most experimental structures have resolved PIEZO channels in an inactive, i.e. curved state. The recently released human structure of PIEZO1 is no exception and shows a symmetrical curvature of all three protein subunits. To achieve a membrane with a curvature complementing the curved protein, multiple approaches were tried (more details can be found in Table S4). In the protocol that we settled on, the first step after preparation of the initial all-atom protein model is the mapping of the protein to a Martini3 CG representation (Fig. 2). Afterwards, the CG model is embedded in a CG, flat membrane consisting of POPC (Fig. 3A). After successful equilibration, while keeping the protein backbone beads restrained, the POPC membrane adapts to the curved membrane and forms a buckle-like shape (Fig. 3B). The membrane starts out flat with a height in one plane at around 100 Å. After a few nanoseconds of simulation, the lipids start to move and indents surrounding the channel blades become visible.

**Fig. 3:**
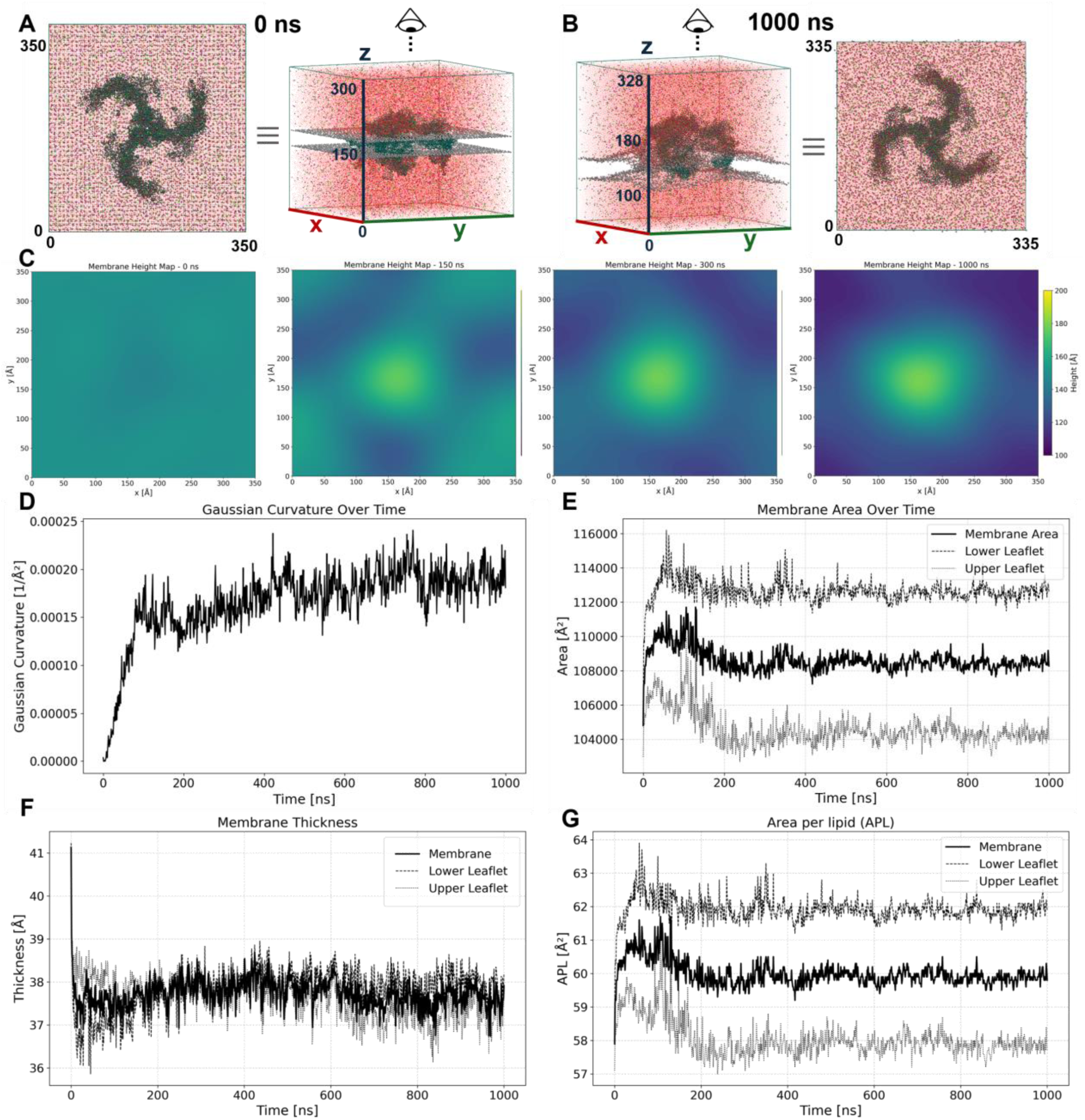
A) Illustration of the simulation box when viewed from above at the start of the CG simulation. During the preparation the protein was oriented with its C-terminal domain (CTD) directed towards the top of the box. B) Same as A) but at the end of the CG simulation. C) The CG system was simulated for 1 µs. The plots of the membrane height display the protein box when viewed from the top, i.e. the intracellular/CTD of the protein is directed towards the top of the box and the Cap domain is pointing towards the bottom (as shown in A and B). At the beginning, the membrane is placed lower than the protein, i.e. closer to the bottom of the box. After a few nanoseconds membrane lipids start to move upwards towards the channel blades, thereby adapting to the protein shape. After 300 ns barely any changes in the membrane height are detectable and the membrane shows a homogeneous curvature around the protein. The pore is approximately in the middle of the simulation box right at the highest membrane point. Towards the edge of the simulation box the membrane flattens out, therefore the shape can be described as buckle-like. Plots depicting time versus D) average Gaussian curvature, E) membrane area, F) membrane thickness and G) APL of the CG simulation are shown. All these values reach an equilibrium approximately after 300 ns.

These lower parts of the membrane progressively get deeper to finally even out, resulting in a homogeneously curved membrane around the three channel subunits (Fig. 3C). This equilibrated system with stable curvature, area, membrane thickness and area-per-lipid (APL) (Figs. 3D-G) was then converted back to all atom, which resulted in a system with 3,797,493 atoms. After successful equilibration of this system, we applied different pressures to simulate the changes in protein structure in response to different pressures.

In a recent study, Vaisey and McKinnon^4^ postulate an activator binding site for PIP2 that enables unbinding of the Latch. As the Latch domain is not included in our model, we cannot investigate these properties here. Therefore, our workflow focuses on the investigation of mechanical activation and deactivation of human PIEZO1.

### PIEZO1 activation involves flattening of both channel and surrounding membrane under pressure

To get a comprehensive and detailed view of the response of the protein to pressure, 1, -1, -5, -10, -20, -30 and -40 bar were initially applied to the inactive system. However, we observed membrane ruptures for most systems with pressures ≤ -20 bar. This was also visible in the changes to the box x-dimensions as the curves showed a sharp decline after a few nanoseconds of simulation (Fig. 4A). We therefore expanded the investigations at lower pressures and added the -3 and -8 bar simulations. The simulations at pressures from 1 to -10 bar were run as five independent replicas. During the simulations, the channel and membrane adopt a flatter shape (Fig. 4B). This is accompanied by an increase in the x and y dimensions of the simulation box while the z dimension shrinks (Fig. 4A, SI Fig. S1). As can be seen in the representative plot of the P_m_10 simulation, the membrane curvature around the protein changes until 35 ns and then plateaus (Fig. 4C). The analysis of the membrane height in the z dimension allows for a more detailed analysis of the membrane shape. At the beginning of each simulation (t = 0 ns), the membrane is homogeneously curved around the curved protein, and the highest point of the membrane is around the protein centre. During the simulation this highest point moves downwards, i.e. the protein moves upwards if viewed from the side. At the end of the P_m_10 simulation, the whole membrane is in one plane (Fig. 4D). The trend observed in the increase in x dimension is also visible for the protein structures. The higher the applied pressure in the simulation, the flatter the protein, i.e. the higher the protein and membrane move in the simulation box (Fig. 4E).

**Fig. 4:**
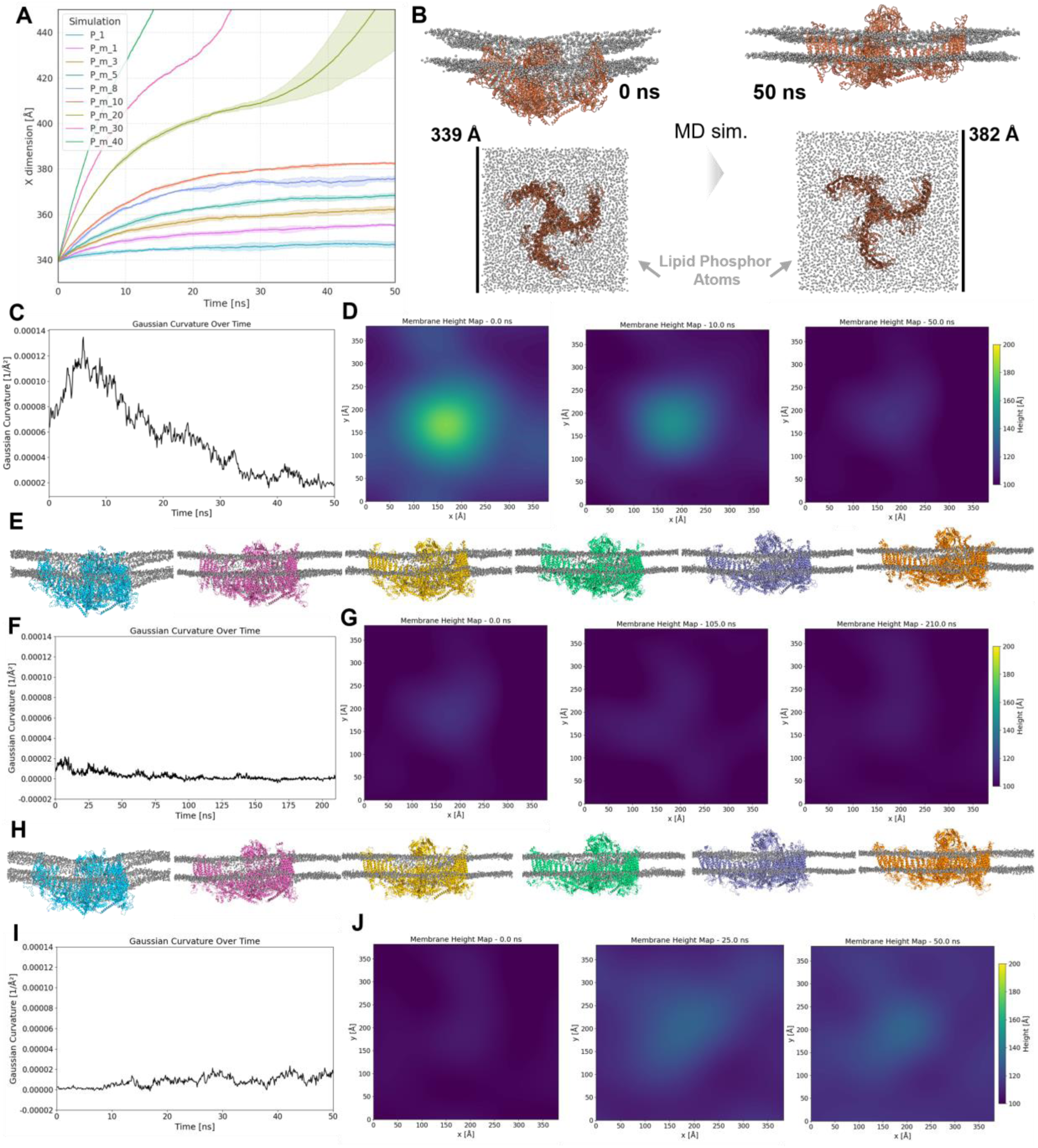
A) Plot describing the simulation time vs. x dimension of the different simulations with different pressure applied. Pressures ≥ 20 bar resulted in membrane ruptures visible as sharp inclines in the plotted lines. In this step all five replicas of the P_1 to P_m_20 were successfully simulated. B) First and last frame of a representative P_m_10 simulation. For a clearer visualisation only lipid phosphor atoms are shown as spheres and the protein is displayed as ribbons. The atoms are coloured grey and the protein is coloured orange. C) Plot of the simulation time versus Gaussian Curvature of the P_m_10 simulation from B). The curve generally flattens out after ∼40 ns. For some simulations the simulations were extended to 100 ns (SI Figs. S8-S26). D) Membrane heights of representative frames from the P_m_10 simulation of B) and C). In the starting structure the membrane is homogeneously curved but starts to flatten out fast. After 10 ns the highest point in the membrane moved down from ∼190 Å to 150 Å. At the end of the simulation almost no difference in membrane height is visible anymore. E) Last frame of a representative replicate of P_1 (blue), P_m_1 (pink), P_m_3 (yellow), P_m_5 (green), P_m_8 (purple) and P_m_10 (orange). Lipid phosphorus atoms are displayed as grey spheres. F) Plot of the To investigate deactivation, the P_m_5 systems and the single stable P_m_10 simulation were subjected to a further simulation step in which pressure coupling was enabled again. The barostat was set to 1 bar resulting in a rapid shrinkage of the membrane, which reached APL, area and thickness values comparable to those of the control P_1 system (SI Figs. S8-S26). The curvature however showed only a low but noticeable increase (Figs. 4I, J).simulation time versus Gaussian Curvature of the P_m_10_cef simulation. G) Membrane heights of representative frames from the P_m_10_cef simulation. H) Same as E) but the last frame of the respective CEF simulation is shown. I) Plot of the simulation time vs. Gaussian Curvature of the P_m_10_1bar simulation. J) Membrane heights of representative frames from the P_m_10_1bar simulation.

The end states of these simulations were taken as starting structures to conduct further 210 ns of simulations where we applied a constant external electric field (CEF) while keeping the volume of the system constant. During these simulations a slight further decrease in curvature is observed (Fig. 4F). This can be interpreted as the further adapting to the flat protein and might be due to the applied electric potential that certainly influences charged protein residues. In the CEF simulations further membrane ruptures were observed for the P_m_8 and P_m_10 simulations, which resulted in the exclusion of two and four replicas, respectively. Therefore, it can be concluded that the P_m_5 and following P_m_5_cef simulations are most suitable for further structural investigations as they allow for sufficient and robust activation.

To further assess the structural response of the membrane and protein under an applied CEF, the curvature and membrane heights were also analysed for a second set of production simulations.

### Pressure-induced channel and membrane flattening modulates ion conductance through pore lipid displacement

We applied a CEF to reproduce the natural transmembrane potential and examine its impact on membrane shape and channel activity. P_m_10_cef shows only a slight further flattening under the simulation conditions (Fig. 4F). For the other simulations a further flattening of the membrane and protein is visible although the NVT ensemble allows for no box size fluctuations. Indeed, the analysis of the membrane apart from curvature shows that it was well equilibrated. For all simulations the APL, area and thickness fluctuated around stable average values (SI Figs. S8-S26A,B,C). However, during the CEF simulations, the curvature around the protein decreased further, accompanied by additional flattening of the protein and the surrounding membrane (Fig. 4G, Figs. S9,11,13,15,17,19,21,23E,F).

Despite the observed curvature change, differences in ion conductance can be observed when inspecting ion permeation events. The experimentally determined conductance of 36 pS is most closely replicated by the P_m_5_cef simulations (Fig. 5A). There is a clear trend observed that the higher the applied pressure, the higher the measured conductance in the simulation (Fig. 5B). Moreover, there is a pronounced trend that the higher the applied pressure in the previous simulation, the larger the distances in the pore are (Fig. 5C). To inspect the resulting trajectories further, we analysed the densities of the lipids and waters surrounding the pore in the centre of the protein. The trend is also reflected in these densities, as in the simulation with the highest pressure applied the lipids completely leave the ion conductance pore while at the same time the water density increases (Fig. 5E). This prompts the idea that lipids are one of the main factors influencing ion conductance in human PIEZO1, which is supported by recent findings from Shan et al.^2^. There, they postulate that lipids seal the pore and are involved in fast inactivation of PIEZO1. This idea is further supported by recent computational and electrophysiological results for mouse PIEZO1 and PIEZO2, where Li et al.^11^ found that channel opening is a two-step mechanism. However, they found similar distances and conductance for mouse PIEZO1 as we did here for the human channel.

**Fig. 5:**
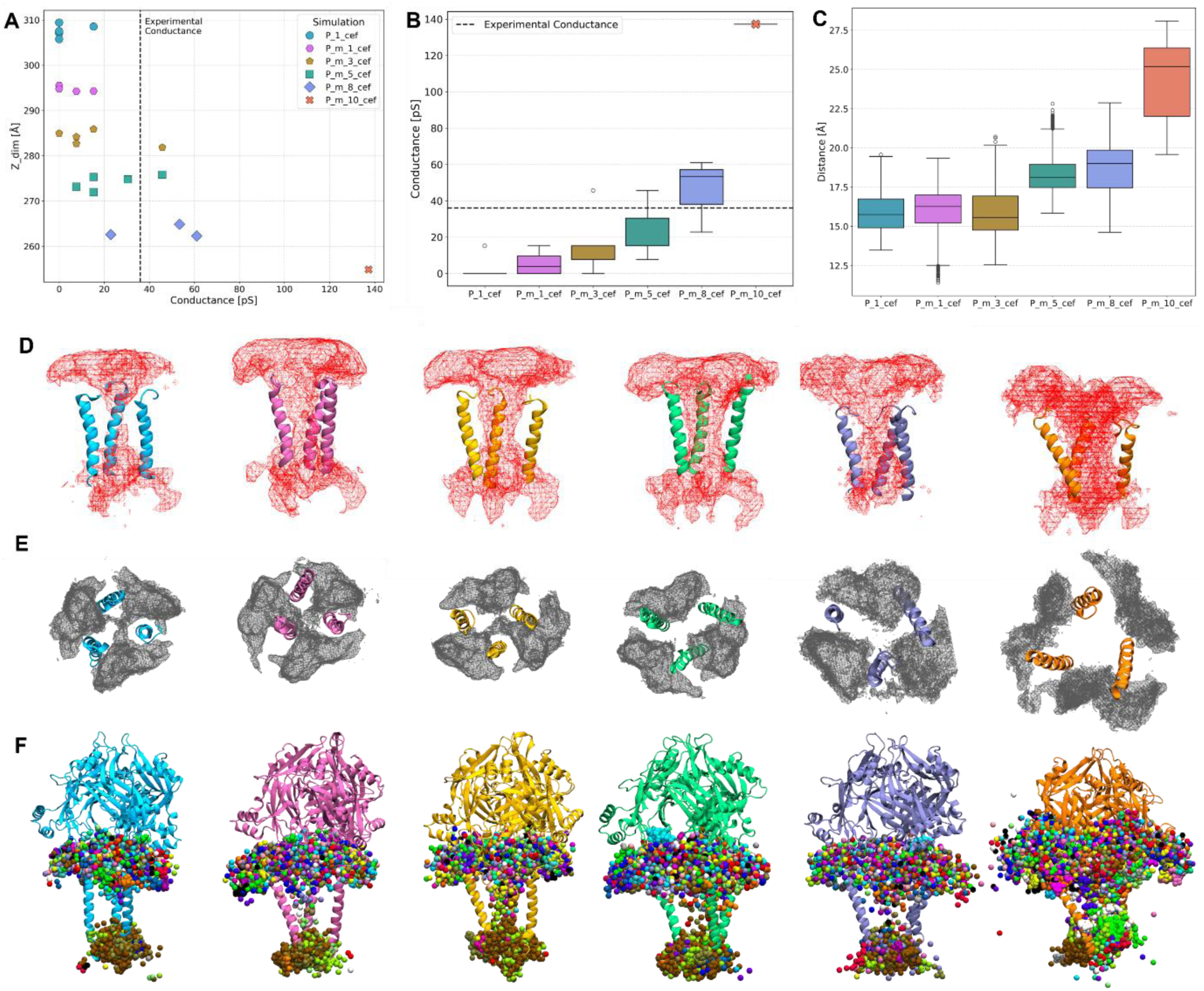
Analysis of the CEF simulations: A) The conductance is plotted against the box z-dimension (Z_dim) of the box. The experimental conductance of PIEZO1 was taken from Gnanasambandam et al.^47^. The smaller the Z_dim, the higher the conductance. B) This can be extended to the pressure, i.e. the higher the applied pressure, the higher the conductance. C) Furthermore, the mean distance of the transmembrane gate also increases the higher the applied pressure. The distance was measured based on the Cβ atom positions of V2450 of all three subunits. D) + E) From left to right: Resulting densities taken from the concatenated trajectories of the P_1_cef, P_m_1_cef, P_m_3_cef, P_m_5_cef, P_m_8_cef and P_m_10_cef simulations for the water (D, red) and lipids (E, grey) within a distance of 15 Å of the pore. F) Same order as before but all second frames of the respective trajectories are shown for the Na^+^ (multi-coloured) within a distance of 15 Å of the pore. The colouring of the protein corresponds to those in the plots A) and B). All data were based on the 200 ns trajectories of five replicas of P_1_cef, P_m_1_cef, P_m_3_cef and P_m_5_cef, three replicas of P_m_8_cef and the single P_m_10_cef replica.

In general, Na^+^ enters the channel in the centre of PIEZO1 from the extracellular side through fenestrations below the Cap domain. Then, it moves through the gap between the three subunits towards the intracellular side and the C-terminal domain (CTD) of the protein. At the bottom of the pore, the ions generally stay for a longer period and move around. Some move upwards again and leave the channel through a similar path as they entered, others can leave the channel through fenestrations at the sides of the pore (Fig. 5F, Movie S1). This behaviour was previously described by Liu et al.^3^ for mouse PIEZO1 and supports their finding that these fenestrations are a restricting factor for ion permeation together with lipids.

To further investigate the role of pore-lining residues in regulating ion permeation and channel structure, we introduced *in silico* mutations at the pore residue R2456 and investigated their effects.

### The R2456-E2117 salt bridge couples pore lipid occupancy to PIEZO1 gating

In recent structural studies, it was proposed that R2456 forms critical salt bridges with the headgroups of pore-occupying lipids keeping them in place and allowing faster inactivation.^2^ The simulation data of wild-type (WT) PIEZO1 also suggest another role of this residue as it forms a very stable salt bridge with E2117. This interaction is conserved in all frames of the simulation, except one replica of P_1_cef (SI Figs. S2,S3). Therefore, this interaction is suggested to be formed independent of the applied pressure and activation state of the channel. E2117 is part of the anchor domain that was shown to be important for mechano-gating. Introduction of more flexibility through mutations in this domain led to i) slower inactivation and ii) made the channel harder to open.^48^ We therefore investigated two mutants that represent these two possibilities, R2456H (GOF) and R2456C (LOF). To investigate the properties of the mutated channels, R2456 of the human PIEZO1 model was *in silico* mutated to either cysteine or histidine. Then, these two systems were simulated at -5 bar, inducing channel and membrane flattening and applied a CEF to promote ion conductance across the membrane. This parameter was chosen as we demonstrated that the conductance of the WT system simulated at this pressure correlates with physiological conductance and all simulated replicas remained stable during simulation.

For the R2456C_cef no ion passage through the channel pore was observed in any of the five replicas (Fig. 6A). The pore remains completely blocked by lipids, contrary to the simulations of WT human PIEZO1 (Figs. 6F,G). This is also evidenced by a lower distance between IHs of the three subunits compared with the WT simulation (Fig. 6C). The cysteine allows for less interactions with E2117 (SI Fig. S3C) and less polar interactions with phospholipid headgroups are possible. Therefore, it is imaginable that the loss of both interactions leads to a less efficient depletion of pore lipids and at the same time the increased flexibility in the anchor domain might impair opening even further (Fig. 6B).

**Fig. 6:**
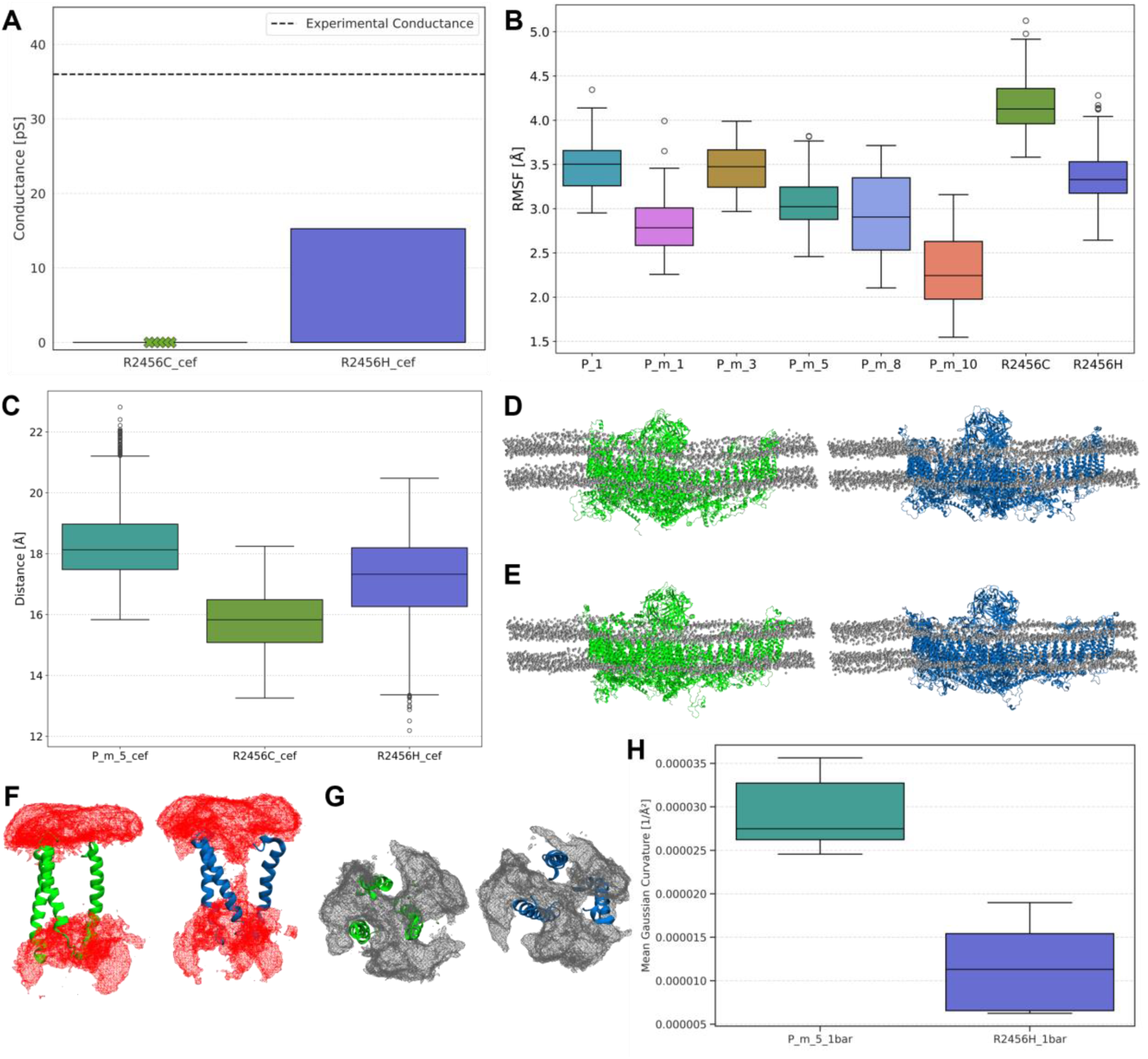
A) Ion conductance of the mutated channel simulations. B) RMSF plots of WT and mutated channel anchor domains (residues 2100 to 2154). C) The mean distance of the transmembrane gate of the mutated channels is lower compared with the WT simulation. The distance was measured based on the Cβ atom positions of V2450 of all three subunits. D) Last frame of a representative replicate of R2456C (green) and R2456H (blue) and E) R2456C_cef (green) and R2456H_cef (blue). F) + G) Resulting densities taken from the concatenated trajectories of R2456C_cef (green) and R2456H_cef (blue) for water (F, red) and lipid (G, grey) molecules within 15 Å of the pore. H) Comparisons of membrane curvature of P_m_5_1bar and R2456H_1bar simulations. The mean values are significantly different (p=0.0062, two-tailed Student’s t-test, T-statistic: 5.44, n=5).

The replicas of R2456H_cef show a decreased ion conductance, which at first might seem surprising since it represents a GOF mutation. It was however shown that the GOF mutation in this case is not the result of an increased ion conductance but a slowed inactivation.^2^ To simulate this, the P_m_5_cef and R2456H_cef systems were subjected to a third simulation where the pressure was set to 1 bar, mimicking the removal of pressure on the system. Indeed, pressure removal induced membrane and channel curvature. However, in the GOF R2456H_1bar system, channel and membrane respond slower compared to P_m_5_1bar system, in line with the reduced inactivation kinetics measured for R2456H *in vitro* compared with the WT channel (Fig. 6H).^49^ This suggests that our simulation protocol not only enables the simulation of activation but also enables the detection of differences in deactivation. Moreover, RMSF plots show the increased flexibility of the anchor domain due to the lack of the stabilising salt bridge (Fig. 6B). The lack of stable interactions, except the typical α-helix backbone interactions with G2452 and S2460, is also visible in the interaction plots of H2456 (SI Fig. S3D).

### Long-range interactions between Cap domain, pore mediate, and THU9 mediate channel activation

The observed differences in ion conductance and membrane shape represent only one aspect of the underlying molecular processes. To move beyond this perspective, we extended our analysis to the WT channel itself, exploring whether differences in its interactions could provide additional insight into the mechanisms driving the observed effects. Therefore, we compared the occurrence of interactions in P_1_cef and P_m_5_cef trajectories, as surrogates for the inactive and active channel states. This analysis revealed an activation mechanism, involving multiple clusters of contacts ranging from the Cap domain to the channel pore.

At the top of the Cap domain T2398 forms less inter-subunit interactions with the neighbouring T2398, i.e. the interactions become less symmetrical. Instead T2398 forms more intra-subunit interactions with N2393, A2395 and L2401. Our results suggest that this change is transmitted downwards towards the pore module, as multiple contacts near the loops connecting the IH/OH and Cap domain (S-linker and OC-linker, respectively) differ between the two channel states. In the inactive P_1_cef trajectory, more interactions were observed between I2440-F2193 and between M2241-M2194. Conversely, upon channel activation, interactions are more tightly formed between the contact pairs G2314-V2427, R2309-G2365, L2432-L2443 and T1992-Q2255. The higher number of contact pairs formed in P_m_5_cef is consistent with Liu et al.^5^ who postulated that S-linker compression is a hallmark of the open-state channel.

The decrease in contact between M2241-M2194 upon activation is accompanied by a concurrent increase in contact between P2001-F1979, indicating that the THU9 helices move closer together in the activated state. Together with M2194, P2001 and F1979 form a hydrophobic cluster that becomes tighter in the activated state (Fig. 7).

**Fig. 7:**
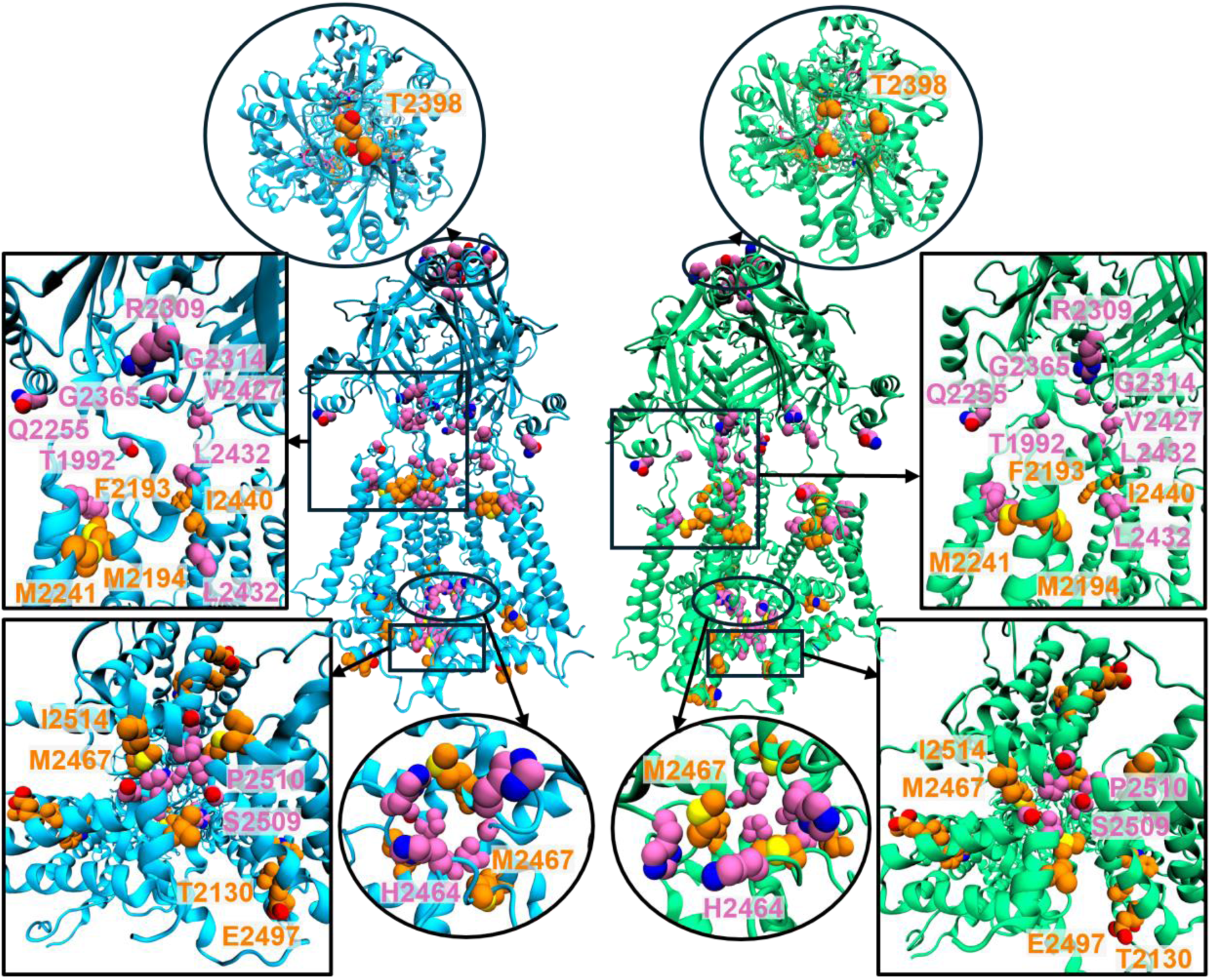
Visualisation of differences in contacts observed between the inactive P_1_cef state (blue) and the active P_m_5_cef state (green). The sidechains of residues mentioned in the text are shown as spheres and labelled accordingly. Contacts were only considered if they differed more than 20% between the simulations and the difference was observed in all three subunits. Contact pairs with more occurrence in P_1_cef are coloured orange and contact pairs with a higher occurrence in P_m_5_cef are coloured pink.

Upon channel activation the following contact changes were observed: The two THU9 helices adjacent to the pore move closer together, as evidenced by tighter contacts between the residue pairs P2001-F1979 and D1968-D2018, while the two distal helices move further outward toward the Blade region. This bending of the two outer THU9 helices towards THU8, results in looser contacts at the top of the helices between V2062-I1974 and I2041-I2010. This also leads to increased contacts at the bottom of the helices between I1952-I1722 and R2028-R1935 in the activated state. Similarly, upon activation these intrahelical unit changes further propagate to the THU8/THU9 interface where, particularly at the top, contacts decrease between three residue pairs: Y2067-I1696, F2046-H1699 and H2042-A1703. At the bottom of THU8, increased contacts are observed between L1800 and W1730 in the activated state (Figs. 8A,B, SI Figs. S4A,B).

**Fig. 8:**
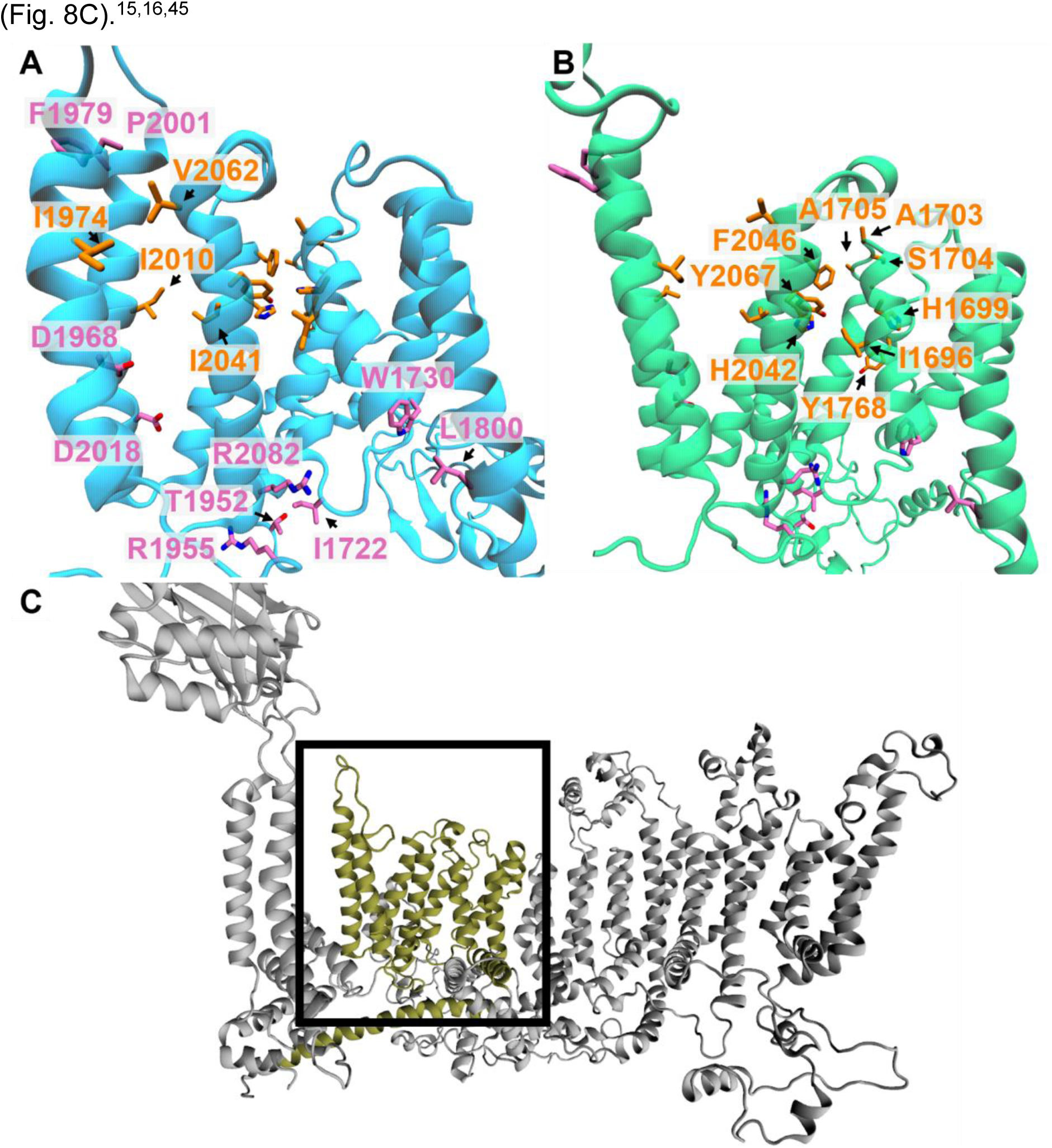
Investigation of the THU8/THU9 interface. Difference in contacts between the A) P_1_cef (blue) and B) P_m_5_cef (green) simulation. Contacts were only considered if they differed more than 20% between the simulations and the difference was observed in all three subunits. Residues with more contacts in P_1_cef, i.e. the non-activated state, are coloured orange. Contacts that are observed more often in P_m_5_cef, i.e. activated state, are coloured pink. C) Location of the THU8/THU9 interface (olive green).

Furthermore, comparison of contacts between P_1_cef and P_m_5_cef simulations suggested an enrichment of activation-associated changes within THU9 and at the THU8/9 interface. Notably, the THU8/THU9 interface corresponds to the proposed yoda1 binding site, which has been validated through multiple mutagenesis studies

### Activation reshapes the yoda1 binding pocket in PIEZO1

Yoda1 is a small-molecule agonist of PIEZO1 that lowers the mechanical threshold for channel activation and stabilises the open state of the channel.^50^ As an additional investigation how interactions around the yoda1 binding site change throughout the simulation, we analysed the P_m_5 trajectories and used the first frame as the reference inactive state, instead of the P_1_cef trajectory. Interestingly, this yielded the additional contact pair W1730-E1688 which is formed during channel activation. These two residues are located directly behind the yoda1 binding pocket in THU8 (Fig. S4C).

Next, we analysed the pocket shapes of the yoda1 binding site for three simulations: the P_1_cef, as surrogate for the inactivated channel state, and the P_m_5_cef and P_m_10_cef simulations, representing two distinct active channel states. This investigation revealed that the THU8/9 interface adopts a narrower and more elongated conformation in the inactive channel (Fig. 9A, SI Figs. S5A,D,G,J). In contrast, in the active state simulations the pocket is more compact, extends further into the THU8/9 interface and is enlarged on the right-hand side along the adjacent helix (Figs. 9B-C, SI Fig. S5).

**Fig. 9:**
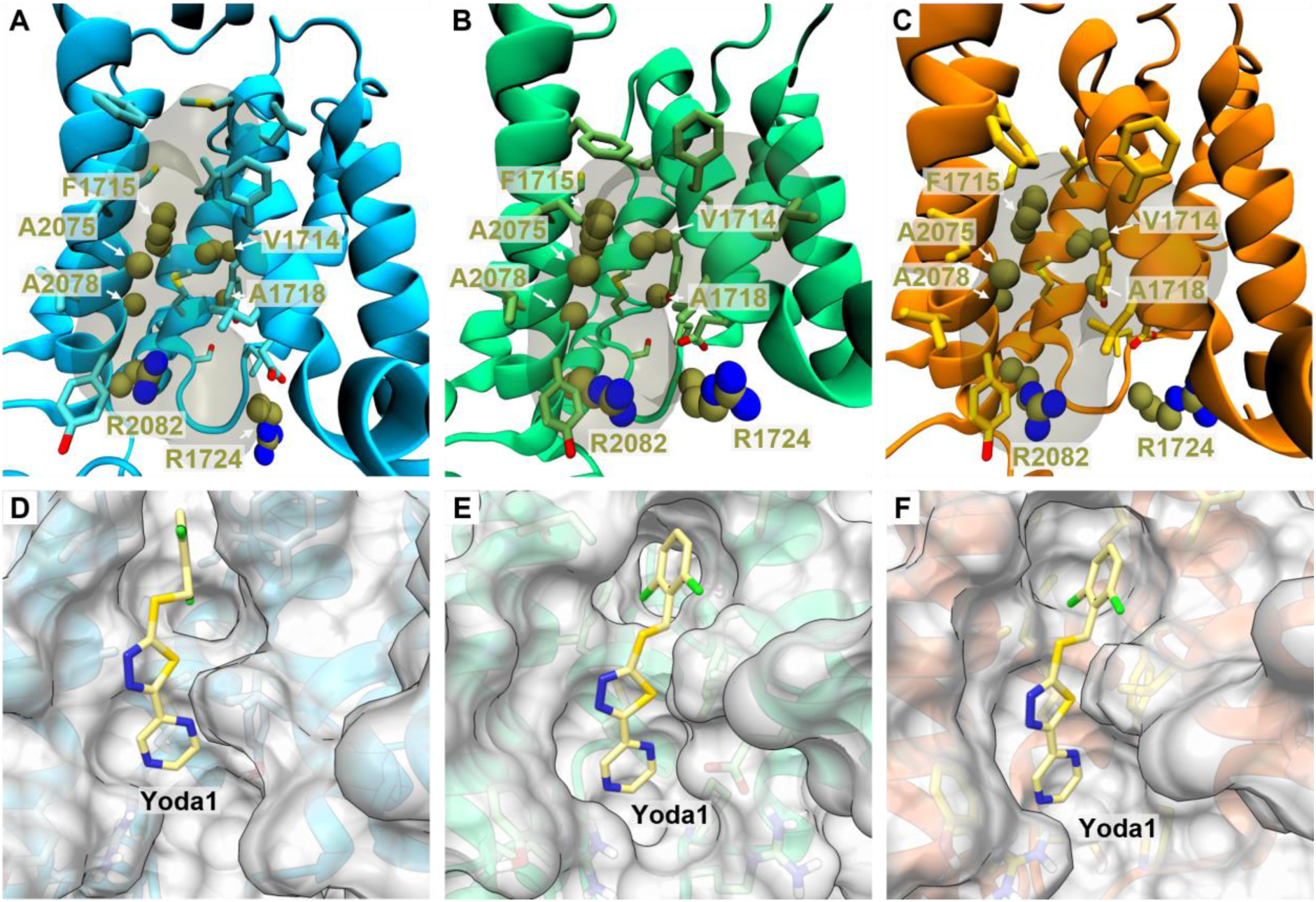
Binding pocket and docking poses of yoda1 at the THU8/9 interface. A), B) C) Pocket shape of yoda1’s binding site as observed in P_1_cef (blue), P_m_5_cef (green) and P_m_10_cef (orange) as detected by fpocket^41^. Residues shown to be involved in yoda1 through mutational studies by Botello-Smith et al.^15^ (A1718, A2075 and A2078), Verkest et al.^16^ (V1714, F1715) and Wijerathne et al.^45^ (R1724 and R2082) are coloured olive green and their heavy atoms are shown as spheres. Binding poses for yoda1 (light yellow) in D) P_1_cef (blue), E) P_m_5_cef (green) and F) P_m_10_cef (orange). The protein surface is shown in light grey for all three simulations.

We used molecular docking to predict the pose of yoda1 to representative channel conformations of P_1_cef (inactive state), P_m_5_cef and P_m_10_cef (active states) (Figs. 9D-F). The binding modes are consistent with the residues identified by Botello-Smith *et al*.^15^ and align with the mutational data of Verkest *et al*.^16^. The near-zero strain of yoda1 in the activated P_m_5_cef pocket (0.5 kcal mol⁻¹), compared with the 3.2 kcal mol⁻¹ penalty in the inactive P_1_cef pocket, therefore increases our confidence that the pose predicted in the activated state reflects a physically plausible, potentially bioactive binding mode.

In detail, the A2078W mutation abolishes yoda1-mediated channel activation^16^, likely by introducing a steric hindrance by the bulky tryptophane side chain. Our docking poses suggest that A2078 is close to the central thiadiazole ring of yoda1 and tryptophane would simply block a large, central area of the pocket. In contrast, the residues F1715 and V1714 are located deeper within the THUs’ interface. The mutations F1715A and V1714A preserved yoda1 binding but impaired its modulation of mechanically evoked currents. These observations are compatible with the model of functionally distinct ligand-interaction regions: a more superficial site that contributes predominantly to activation and a deeper site involved in the modulation of blade curvature and mechanotransduction. These results are further supported by free-energy calculations by Jiang et al.^14^. We therefore propose that channel activation is associated with reshaping of the THU8/THU9 pocket, thereby favouring yoda1 binding, which in turn then stabilises the activated conformation of PIEZO1.

Within this framework, F1715 may be particularly important for transmitting ligand-induced conformational changes rather than for ligand binding per se, whereas A2078 may contribute more directly to the accessibility and geometry of the binding pocket. In our simulations containing a membrane environment, M1719 does not obstruct the proposed site in contrast to other structural models^16^, which further supports the notion that yoda1 recognition depends on the conformational and membrane-dependent dynamics of this region. Recent evidence identifying additional contributions from arginine residues within the same binding region provides further support for a structurally and functionally heterogeneous yoda1-binding interface.^45^

### The Beam domain undergoes a lever-like rearrangement during PIEZO1 activation

Below the THU8/THU9 interface there is the Beam domain which shows another notable difference between the activated, i.e. P_m_5_cef, and inactive, i.e. P_1_cef, simulation. On a superficial level, the beam moves further upwards towards the THU interface between its N-terminal end and its C-terminal end below the pore. This not only changes its position in space but changes the angle and orientation between the three helices found at its N-terminal end (SI Figs. S6A,B). The analysis of contact differences shows four notable contact pairs. If viewed from the side, the pairs in front of the Beam, L1319-L1565 and R1323-L1568 form more interactions and move closer together in the active state. Directly at the nick of the Beam L1320 is found in P_1_cef. Upon activation, it forms less contacts with the clasp residue L1543 which is situated behind the Beam in THU8. At the C-terminal end of the Beam, its residue N1361 and the CTD residue L2498 form looser interactions in the activated state (Fig. 10).

**Fig. 10:**
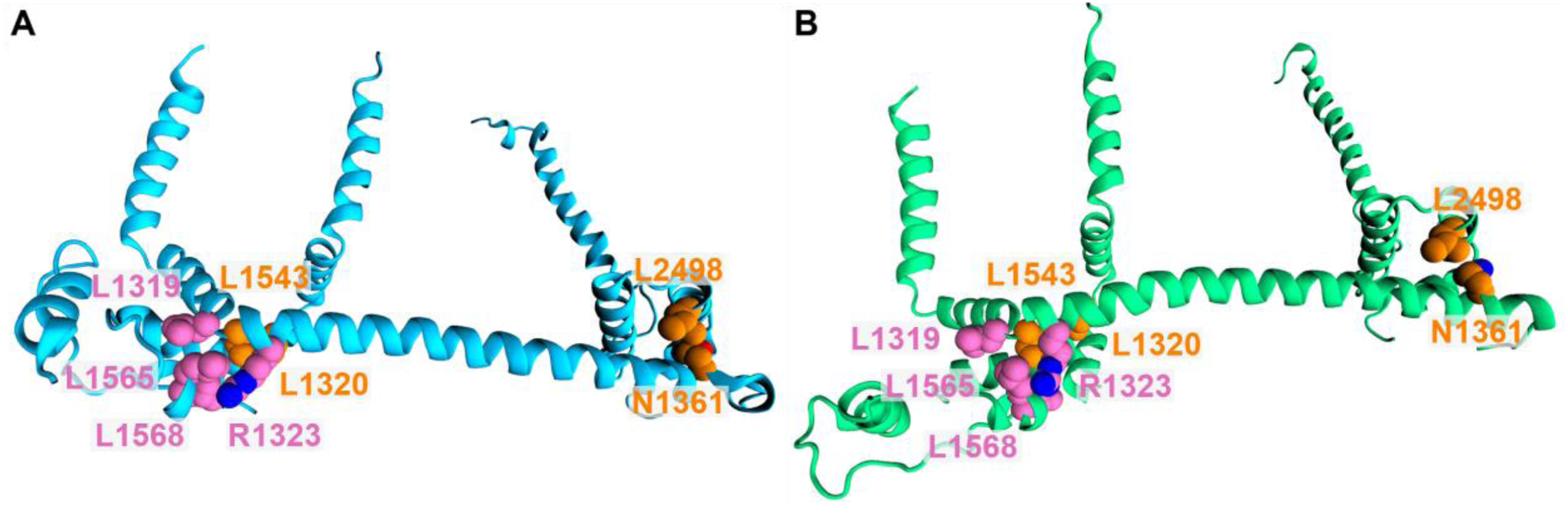
Notable contact pair changes in the beam and clasp domain. Contacts were only considered if they differed more than 20% between the simulations and the difference was observed in all three subunits. Residues that are lost upon activation are coloured orange and contacts that are observed more often in the P_m_5_cef, i.e. active state, are coloured pink.

Additionally, further along the Blades, activation-associated in changes were also observed between residue pairs within THU4-THU7. As these regions include residues that were modelled rather than resolved in the experimental, we did not analyse them in further detail. However, the overall pattern can be summarised as tighter contacts at the bottom and looser contacts at the top of these THUs during channel activation, mirroring the trend observed for THU8/9 (SI Figs. S6C,D).

## Conclusion

We combined CG and atomistic MD simulations under controlled membrane tension and an applied transmembrane electric field to reconstruct the full activation-deactivation cycle of human PIEZO1. This provides, to our knowledge, the first mechanistic view of the human PIEZO1 gating cycle at atomic resolution and offers a structural rationale for (dys)function-associated mutations of the channel.

Systematic pressure adaptation identified a robust activation regime (P_m_5) in which the channel and surrounding membrane flattens and its conductance reproduces the experimental value of 36 pS. Our simulations suggest that ion permeation is gated by the displacement of pore lipids, providing atomistic support for the lipid-sealing model of PIEZO1 inactivation. *In silico* mutagenesis of R2456 suggests that the salt bridge with E2117 in the anchor domain linked to the pore lipid displacement and disrupting this interaction reproduced the gain- and loss-of-function phenotypes of R2456H and R2456C, respectively.

Furthermore, our simulations of human PIEZO1 activation and deactivation suggest rearrangements of interactions that link the Cap domain, the S- and OC-linkers, a hydrophobic THU9 cluster, in addition to a lever-like reorientation of the Beam domain, and a remodelling of the THU8/THU9 yoda1 binding pocket. As to the latter, activation-dependent reshaping of the yoda1 pocket could explain the state-dependent action of yoda1 and provides the structural basis for the design of next-generation PIEZO1 modulators.

## Supporting information

p_1_cef_inactive-state_representative-docking-pose-yoda1

p_m_5_cef_active-state_representative-docking-pose-yoda1

## Acknowledgements

This work was supported by the Research Training Group “Chemical biology of ion channels (Chembion)” funded by the *Deutsche Forschungsgemeinschaft* (DFG), which is gratefully acknowledged. O.K. was funded by a Heisenberg-Professorship (DFG; KO 4689/5-1 & KO 4689/5-2). J.M. and O.K. gratefully acknowledge the HPC resources and support provided by the Erlangen National High Performance Computing Centre (NHR@FAU) of the Friedrich-Alexander-Universität Erlangen-Nürnberg (FAU) under the NHR project k101ee. NHR funding is provided by federal and Bavarian state authorities.

