## Supporting Information for "The activation-deactivation cycle of human PIEZO1"

**Table S1:** Overview of the final all-atom composition of the three prepared systems.

| System | Total No. of atoms | No. of protein atoms | No. of $Na^+$ | No. of $Cl^-$ | No. of Water Molecules | No. of Lipids (upper leaflet) | No. of Lipids (lower leaflet) | Box size after equilibration [Å$^3$] |
|---|---|---|---|---|---|---|---|---|
| **Wild type** | 3,797,493 | 88,372 | 2,847 | 2,847 | 1,072,740 | 1,802 | 1,819 | 339 x 339 x 323 |
| **R2456C** | 3,797,591 | 88,342 | 2,850 | 2,847 | 1,072,740 | 1,802 | 1,820 | 339 x 339 x 324 |
| **R2456H** | 3,797,475 | 88,351 | 2,850 | 2,847 | 1,072,740 | 1,802 | 1,819 | 338 x 338 x 325 |

**Table S2:** Box sizes of the systems simulated with an applied constant external electric field (cef).

| System | Replica | x-dimension [Å] | y-dimension [Å] | z-dimension [Å] |
|---|---|---|---|---|
| **P_1_cef** | 1 | 348.0 | 348.0 | 307.1 |
| | 2 | 347.7 | 347.7 | 307.5 |
| | 3 | 346.7 | 346.7 | 309.4 |
| | 4 | 347.2 | 347.2 | 308.5 |
| | 5 | 348.8 | 348.8 | 305.7 |
| **P_m_1_cef** | 1 | 354.9 | 354.9 | 295.5 |
| | 2 | 355.5 | 355.5 | 294.3 |
| | 3 | 355.5 | 355.5 | 294.3 |
| | 4 | 355.1 | 355.1 | 295.0 |
| | 5 | 355.2 | 355.2 | 294.8 |
| **P_m_3_cef** | 1 | 360.7 | 360.7 | 285.9 |
| | 2 | 361.3 | 361.1 | 285.0 |
| | 3 | 363.7 | 363.7 | 281.2 |
| | 4 | 362.8 | 362.8 | 282.7 |
| | 5 | 361.7 | 361.7 | 284.2 |
| **P_m_5_cef** | 1 | 369.8 | 369.8 | 272.0 |
| | 2 | 369.0 | 369.0 | 273.2 |
| | 3 | 367.7 | 367.7 | 275.3 |
| | 4 | 367.9 | 367.9 | 274.8 |
| | 5 | 367.2 | 367.2 | 275.8 |
| **P_m_8_cef** | 1 | 376.4 | 376.4 | 262.6 |
| | 2 | 374.9 | 374.9 | 264.9 |
| | 3 | 376.6 | 376.6 | 262.3 |
| **P_m_10_cef** | 1 | 382.1 | 382.1 | 254.9 |
| **R2456C_cef** | 1 | 365.9 | 365.9 | 278.0 |
| | 2 | 366.8 | 366.8 | 276.6 |
| | 3 | 368.7 | 368.7 | 273.7 |
| | 4 | 367.8 | 367.8 | 275.0 |
| | 5 | 366.7 | 366.7 | 276.7 |
| **R2456H_cef** | 1 | 367.7 | 367.7 | 275.4 |
| | 2 | 369.2 | 369.2 | 272.8 |
| | 3 | 370.7 | 370.8 | 270.6 |
| | 4 | 370.3 | 370.3 | 271.3 |
| | 5 | 368.2 | 368.2 | 274.5 |

**Table S3:** Overview of individual conductance of the simulated systems.

| System | Replica | Voltage [V] | No. of crossings | Current [pA] | Conductance [pS] |
|---|---|---|---|---|---|
| **P_1_cef** | 1 | 0.1 | 0 | 0 | 0 |
| | 2 | 0.1 | 0 | 0 | 0 |
| | 3 | 0.1 | 0 | 0 | 0 |
| | 4 | 0.1 | 1 | 1.53 | 15.26 |
| | 5 | 0.1 | 0 | 0 | 0 |
| **P_m_1_cef** | 1 | 0.1 | 0 | 0 | 0 |
| | 2 | 0.1 | 2 | 1.53 | 15.26 |
| | 3 | 0.1 | 1 | 0.76 | 7.63 |
| | 4 | 0.1 | 1 | 0.76 | 7.63 |
| | 5 | 0.1 | 0 | 0 | 0 |
| **P_m_3_cef** | 1 | 0.1 | 2 | 1.53 | 15.26 |
| | 2 | 0.1 | 0 | 0 | 0 |
| | 3 | 0.1 | 6 | 4.58 | 45.77 |
| | 4 | 0.1 | 1 | 0.76 | 7.63 |
| | 5 | 0.1 | 1 | 0.76 | 7.63 |
| **P_m_5_cef** | 1 | 0.1 | 2 | 1.53 | 15.26 |
| | 2 | 0.1 | 1 | 0.76 | 7.63 |
| | 3 | 0.1 | 2 | 1.53 | 15.26 |
| | 4 | 0.1 | 4 | 3.05 | 30.51 |
| | 5 | 0.1 | 6 | 4.58 | 45.77 |
| **P_m_8_cef** | 1 | 0.1 | 3 | 2.29 | 22.89 |
| | 2 | 0.1 | 7 | 5.34 | 53.40 |
| | 3 | 0.1 | 2 | 1.53 | 15.26 |
| **P_m_10_cef** | 1 | 0.1 | 18 | 13.7 | 137.31 |
| **R2456C_cef** | 1 | 0.1 | 0 | 0 | 0 |
| | 2 | 0.1 | 0 | 0 | 0 |
| | 3 | 0.1 | 0 | 0 | 0 |
| | 4 | 0.1 | 0 | 0 | 0 |
| | 5 | 0.1 | 0 | 0 | 0 |
| **R2456H_cef** | 1 | 0.1 | 0 | 0 | 0 |
| | 2 | 0.1 | 0 | 0 | 0 |
| | 3 | 0.1 | 1 | 1.53 | 15.26 |
| | 4 | 0.1 | 1 | 1.53 | 15.26 |
| | 5 | 0.1 | 0 | 0 | 0 |

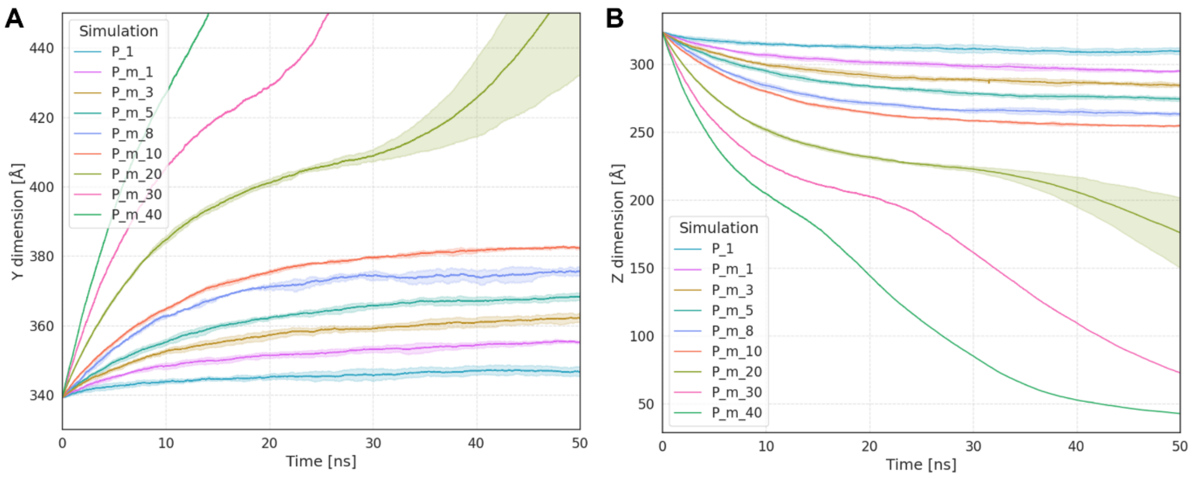


**Fig. S1:** Illustrations of the simulation time vs. A) y dimension and B) vs. z dimension of the eight different simulations each with a different applied pressure.

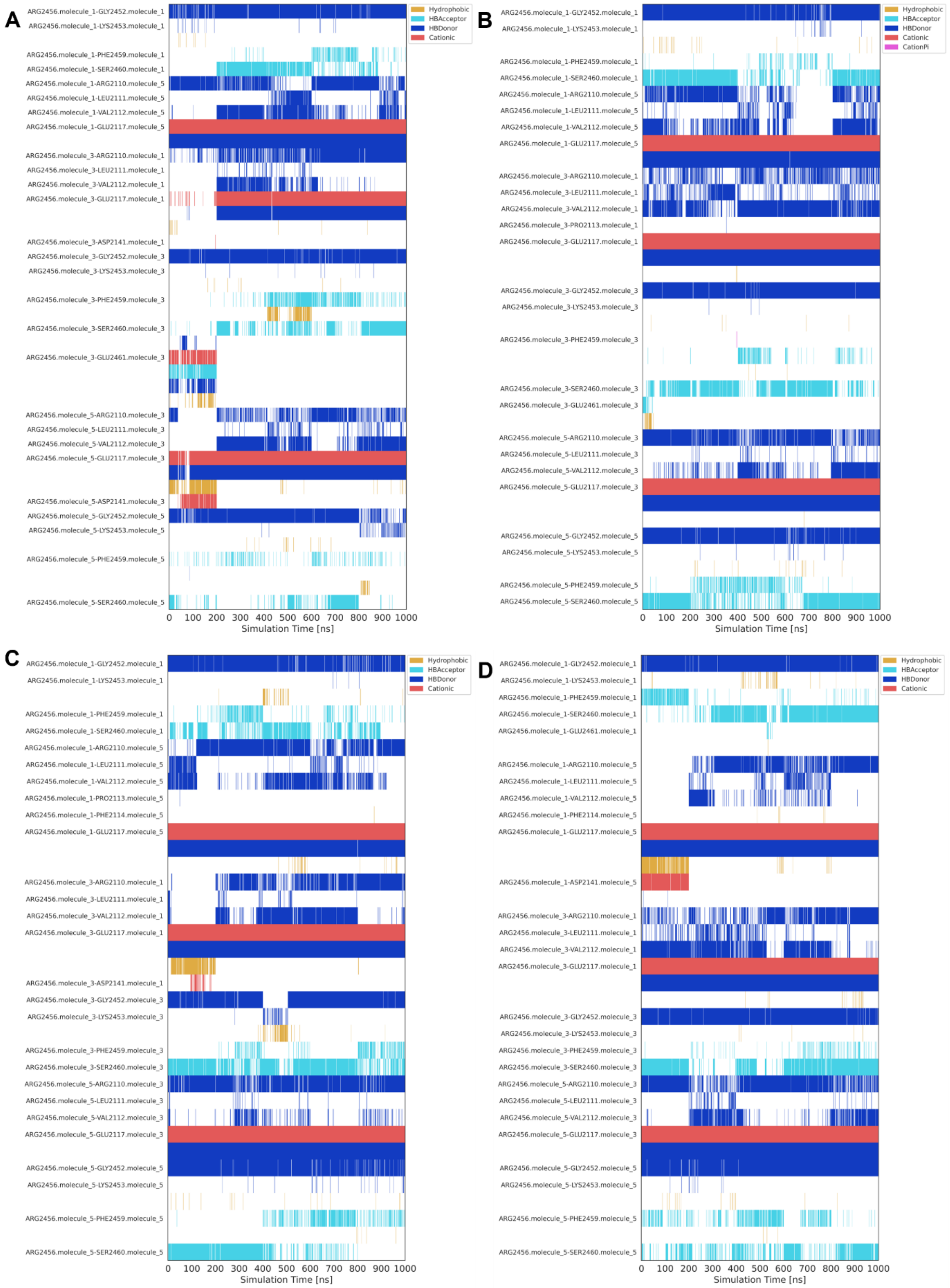


**Fig. S2:** Interaction of R2456 in A) P_1_cef, B) P_m_1_cef, C) P_m_3_cef and D) P_m_5_cef as detected by ProLIF.[1]

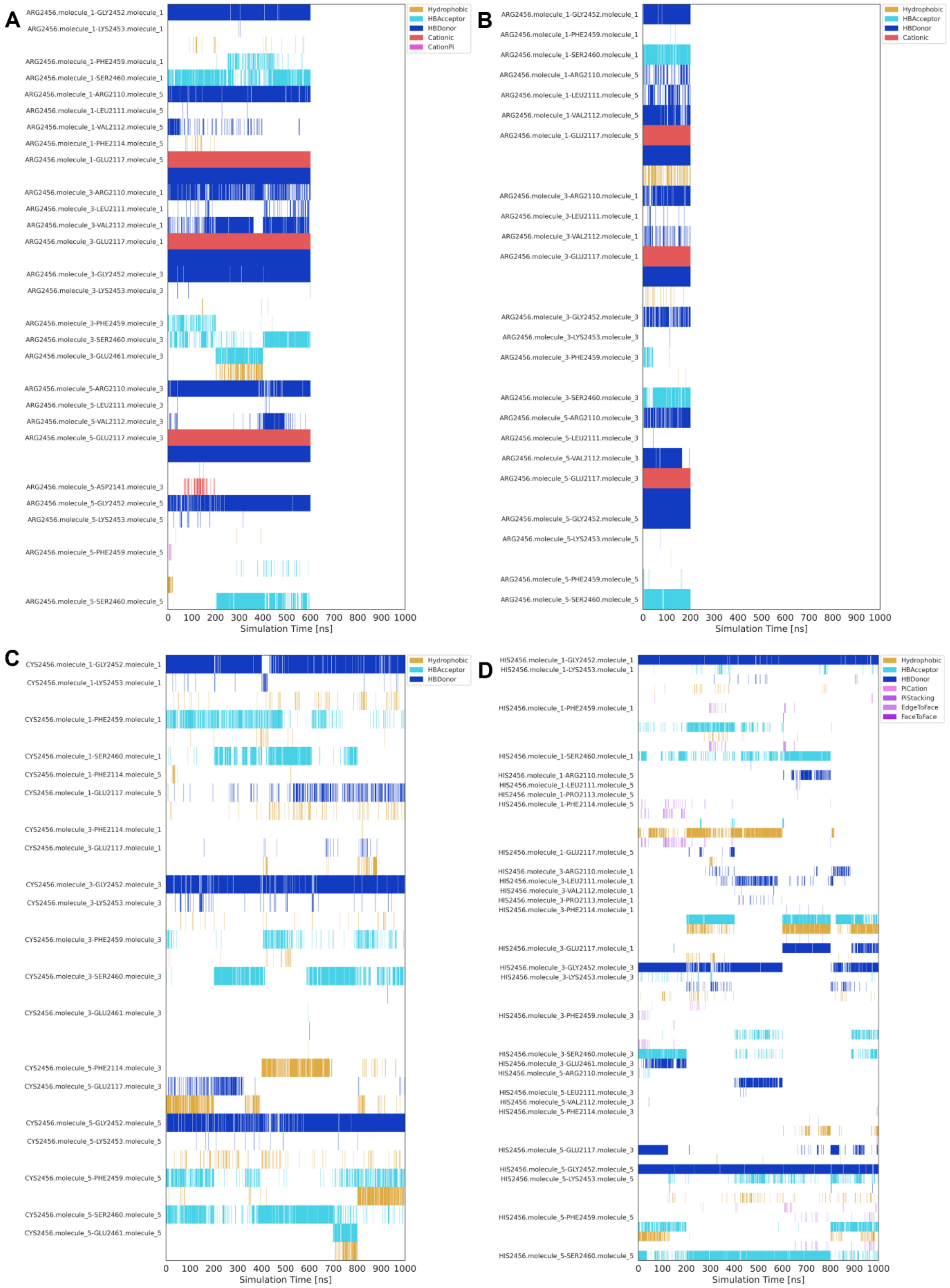


**Fig. S3:** Interaction of R2456 in A) P_m_8_cef, B) P_m_10_cef. C) Interaction of C2456 in R2456C_cef and D) H2456 in R2456H_cef as detected by ProLIF.[1]

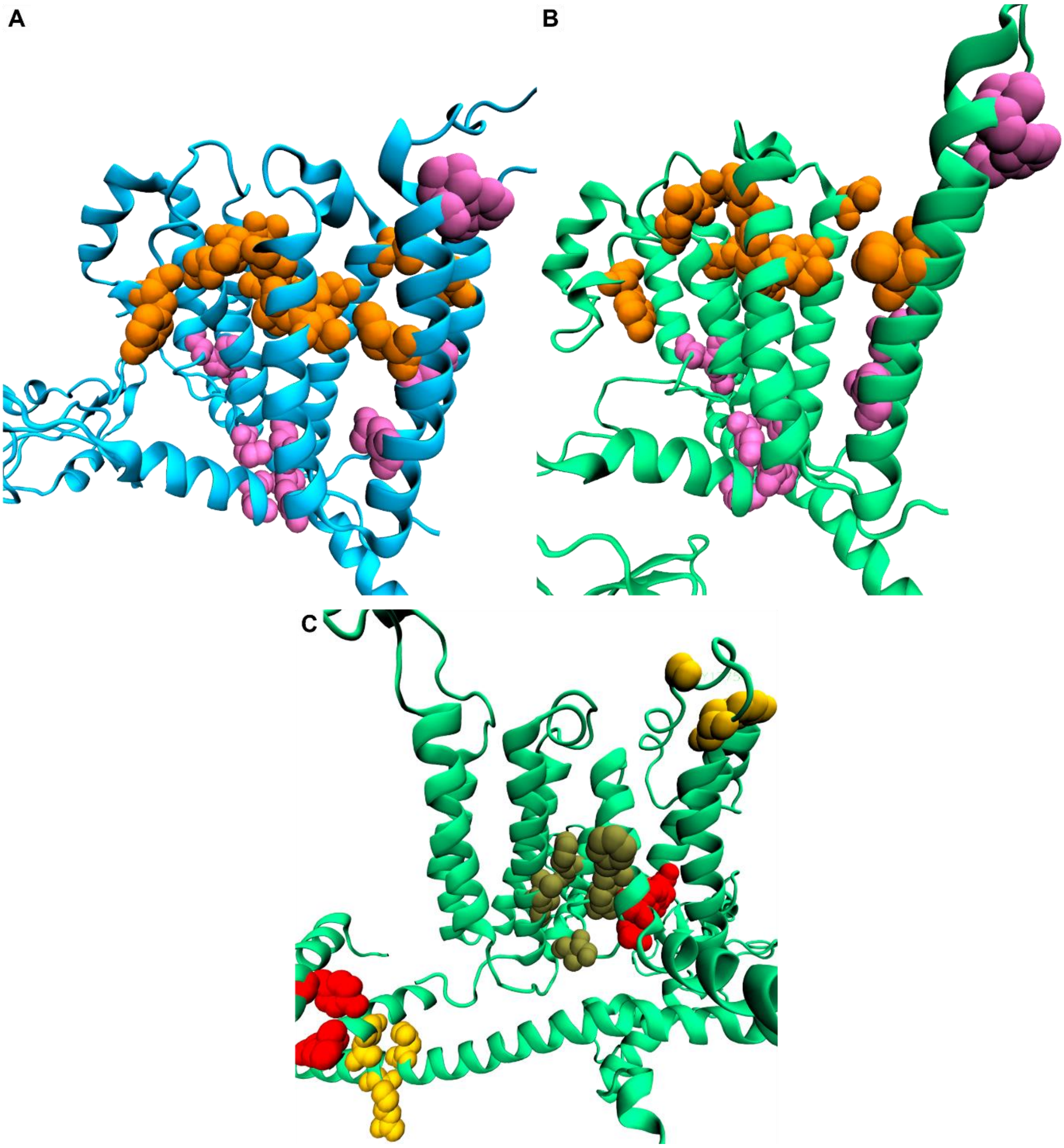


**Fig. S4:** A) Top changed interactions in P_1_cef (blue) and B) P_m_5_cef (green). Contacts were only considered if they differed more than 20% between the simulations and the difference was observed in all three subunits. Residues with more contacts in P_1_cef, i.e. non-activated state, are coloured orange. Contacts that are observed more often in P_m_5_cef, i.e. activated state, are coloured pink. C) Top changed interaction pairs if the first frame of P_m_5_cef is compared with the rest of the trajectory. Residues of the yoda1 binding site[2–4] are shown as olive coloured spheres. Interaction pairs detected in the starting frame and less in the trajectory are coloured yellow, while the ones missing in the starting frame are coloured red, i.e. they are formed in all three subunits during the simulation.

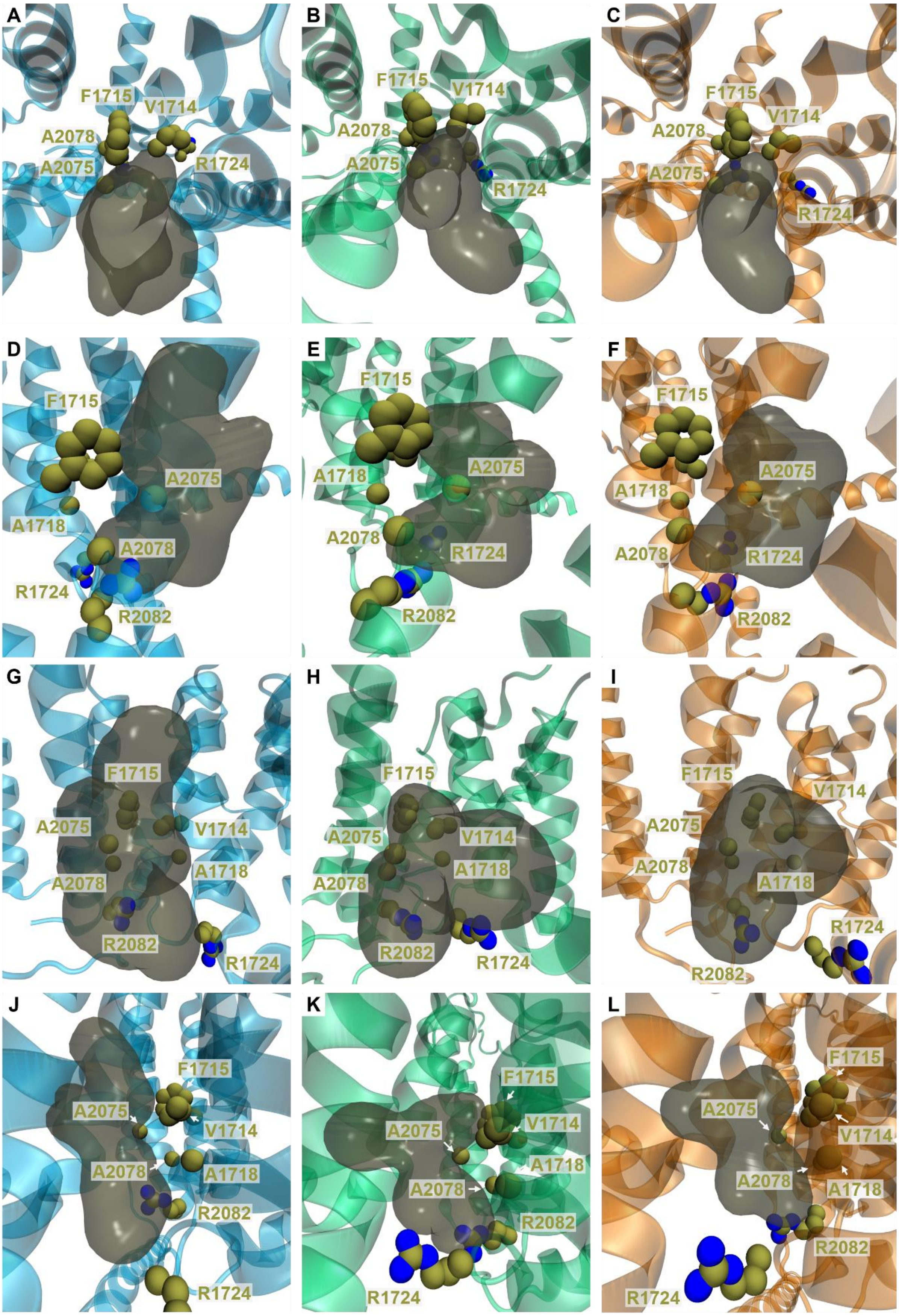


**Fig. S5:** Pocket shapes for the yoda1 pockets of P_1_cef (blue), P_m_5_cef (green) and P_m_10_cef (orange) as identified by fpocket[5] when viewed from the top (A, B, C), left side (D, E, F), front (G, H, I) and right side (J, K, L).

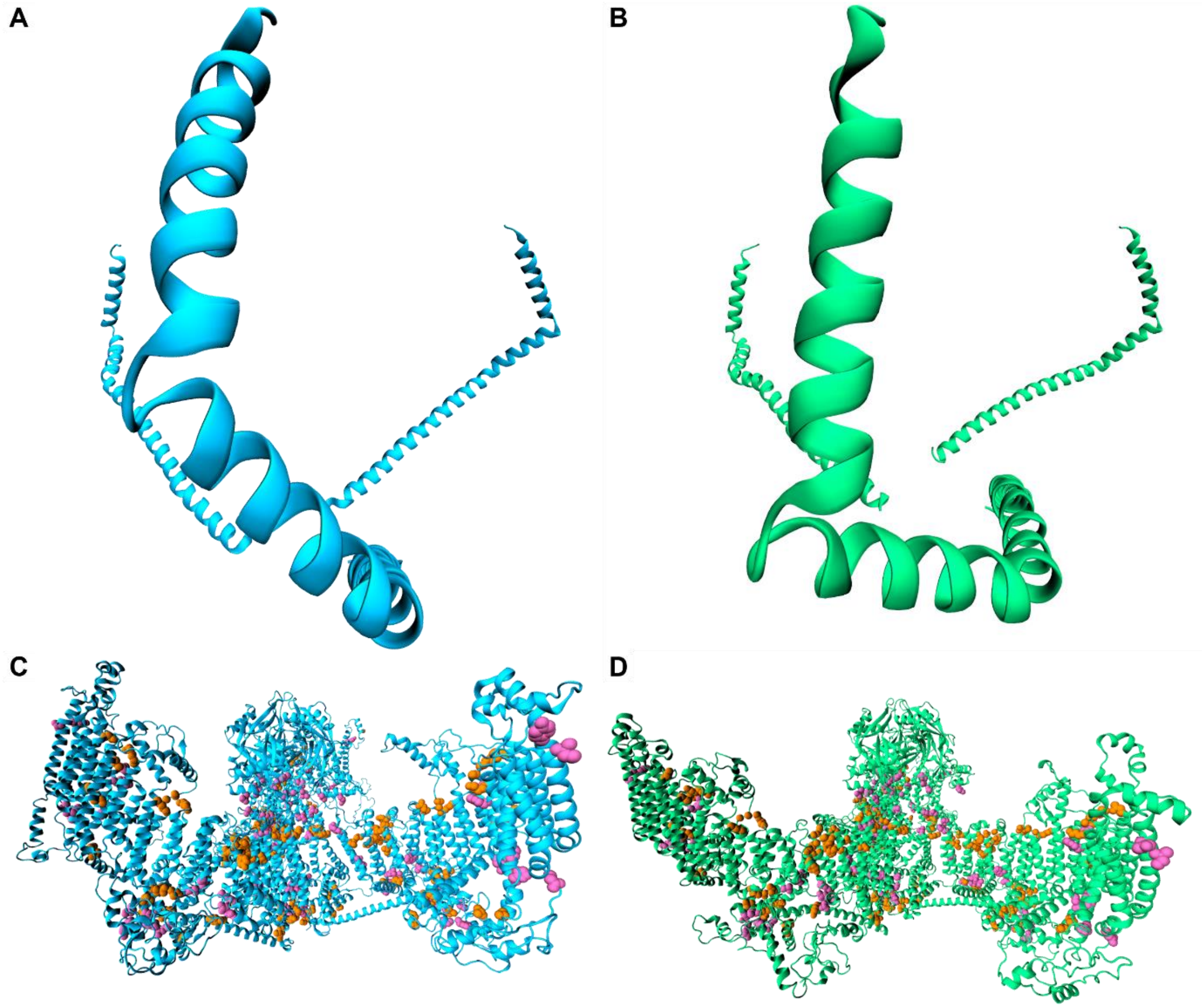


**Fig. S6:** Beam shape of the last frame of A) P_1_cef (blue) and B) P_m_5_cef (green). C) All differences in contact pairs for C) P_1_cef and D) P_m_5_cef. Contacts were only considered if they differed more than 20% between the simulations and the difference was observed in all three subunits. Residues with more contacts in P_1_cef, i.e. non-activated state, are coloured orange. Contacts that are observed more often in P_m_5_cef, i.e. activated state, are coloured pink.

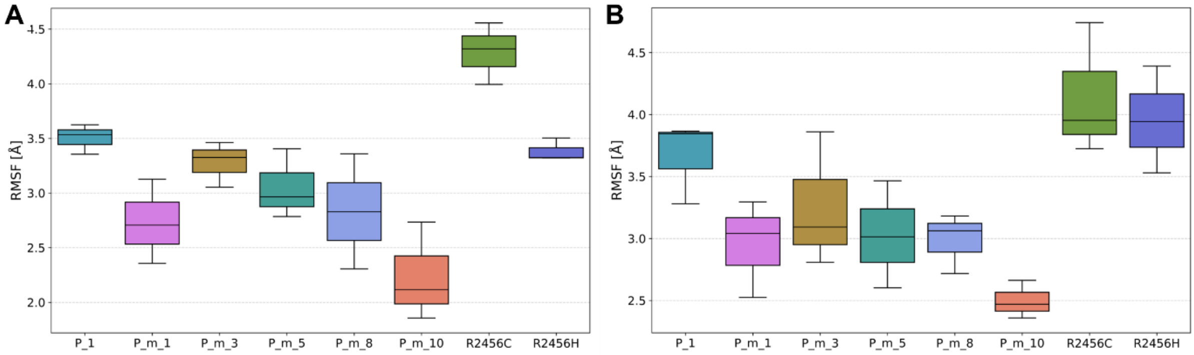


**Fig. S7:** RMSF plots of A) E2117 and B) R2456 or its corresponding mutated amino acid for R2456C_cef (C2456) and R2456H_cef (H2456).

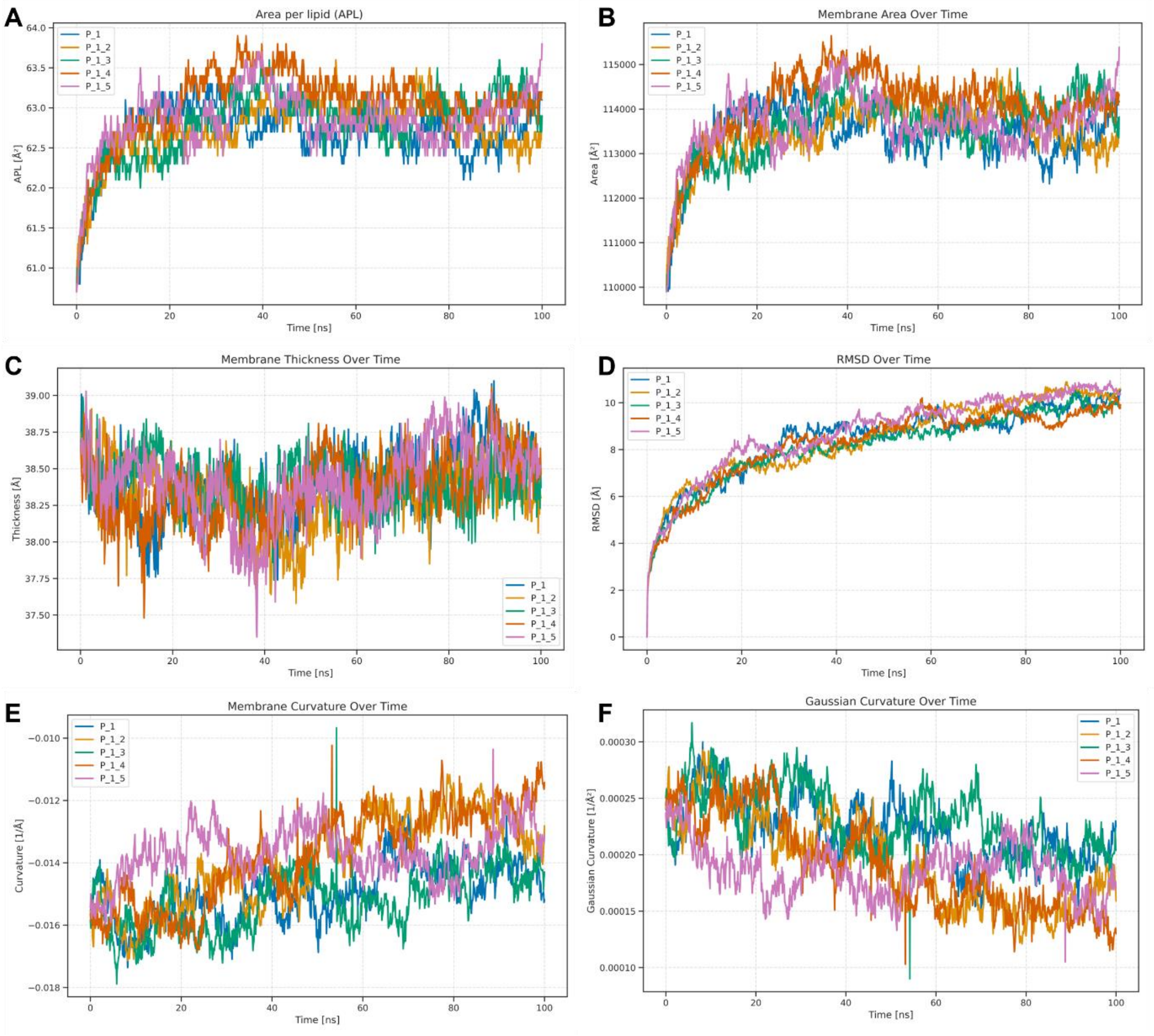


**Fig. S8:** Properties of P_1.

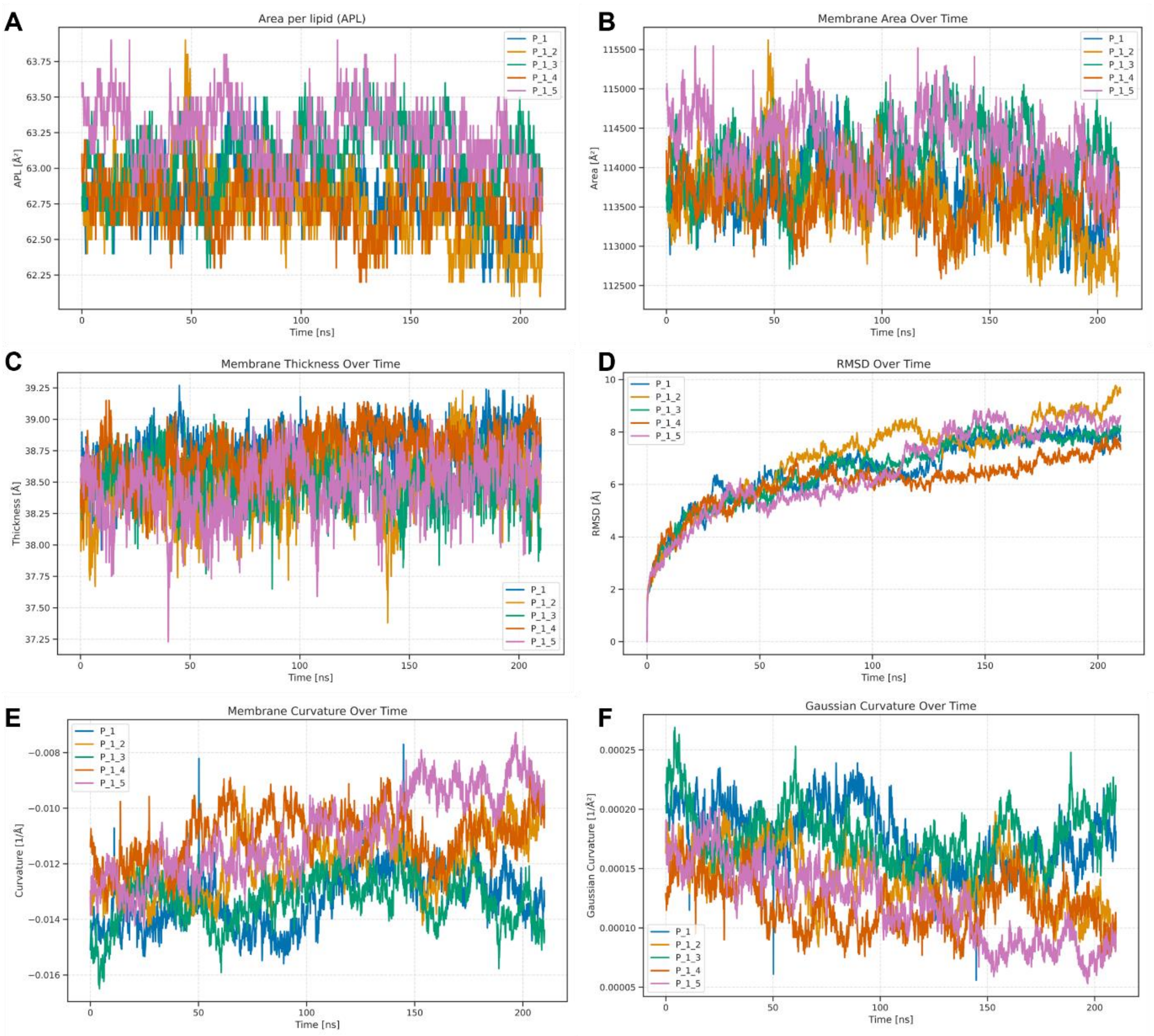


**Fig. S9:** Properties of P_1_cef.

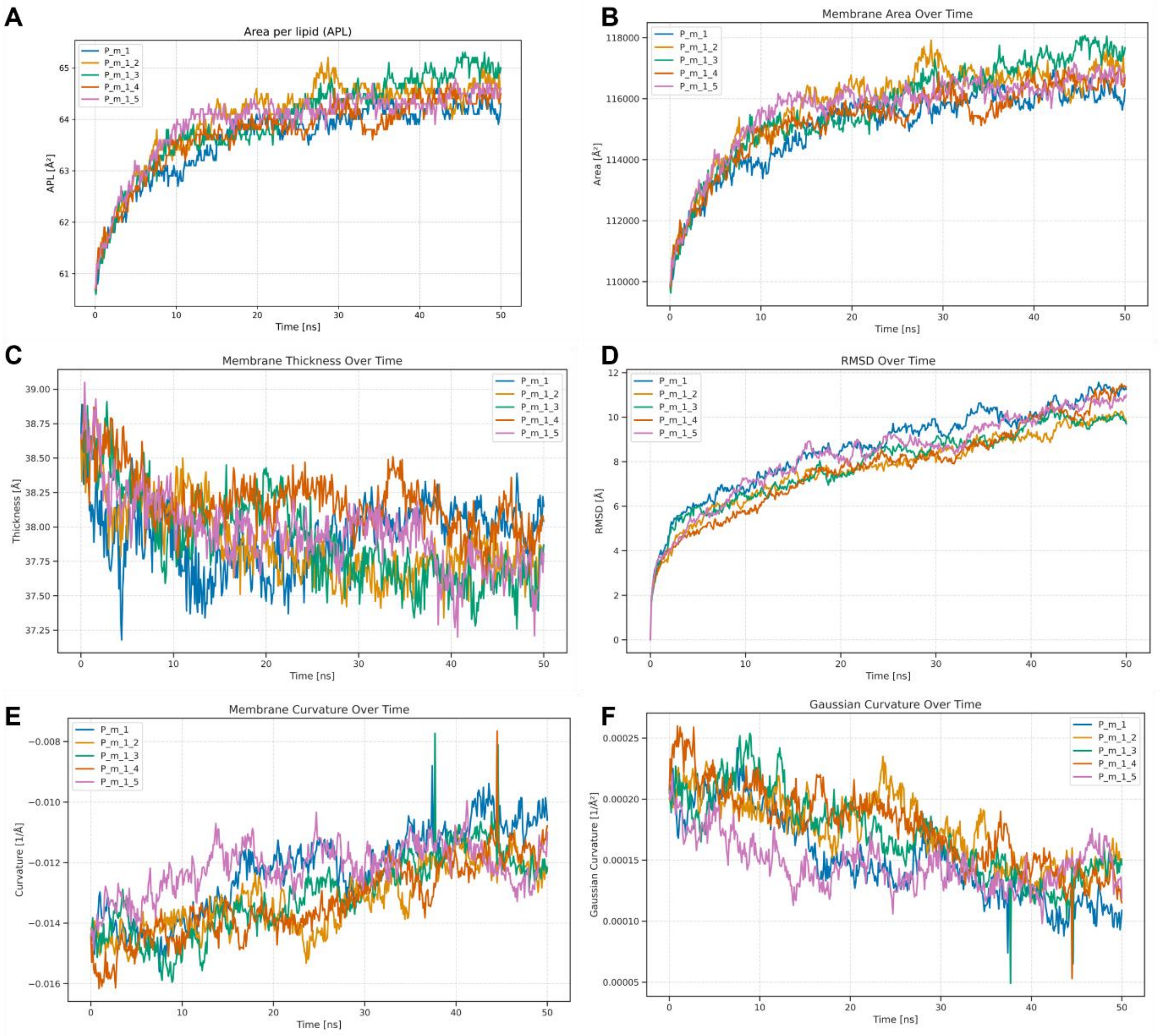


**Fig. S10:** Properties of P_m_1.

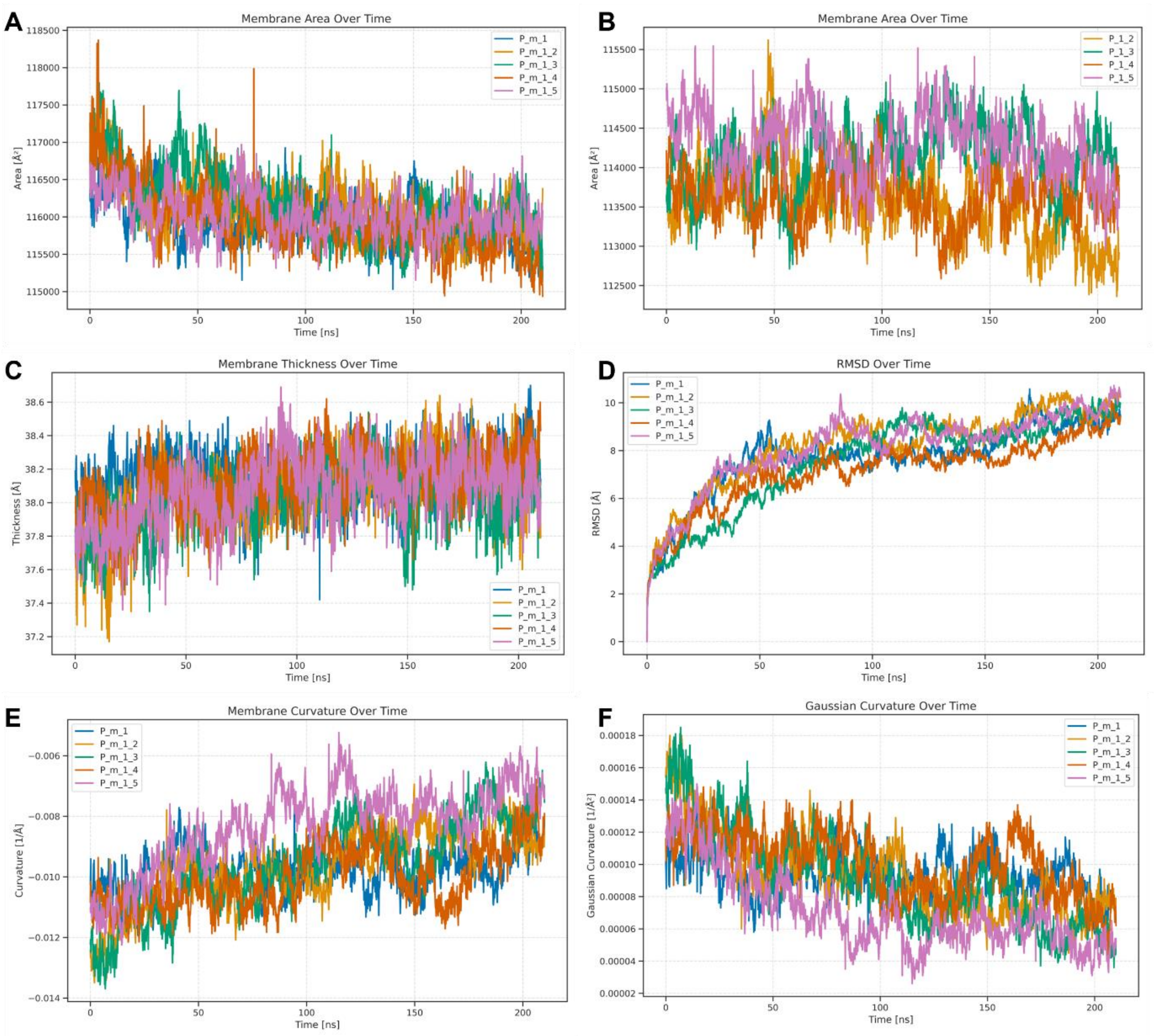


**Fig. S11:** Properties of P_m_1_cef.

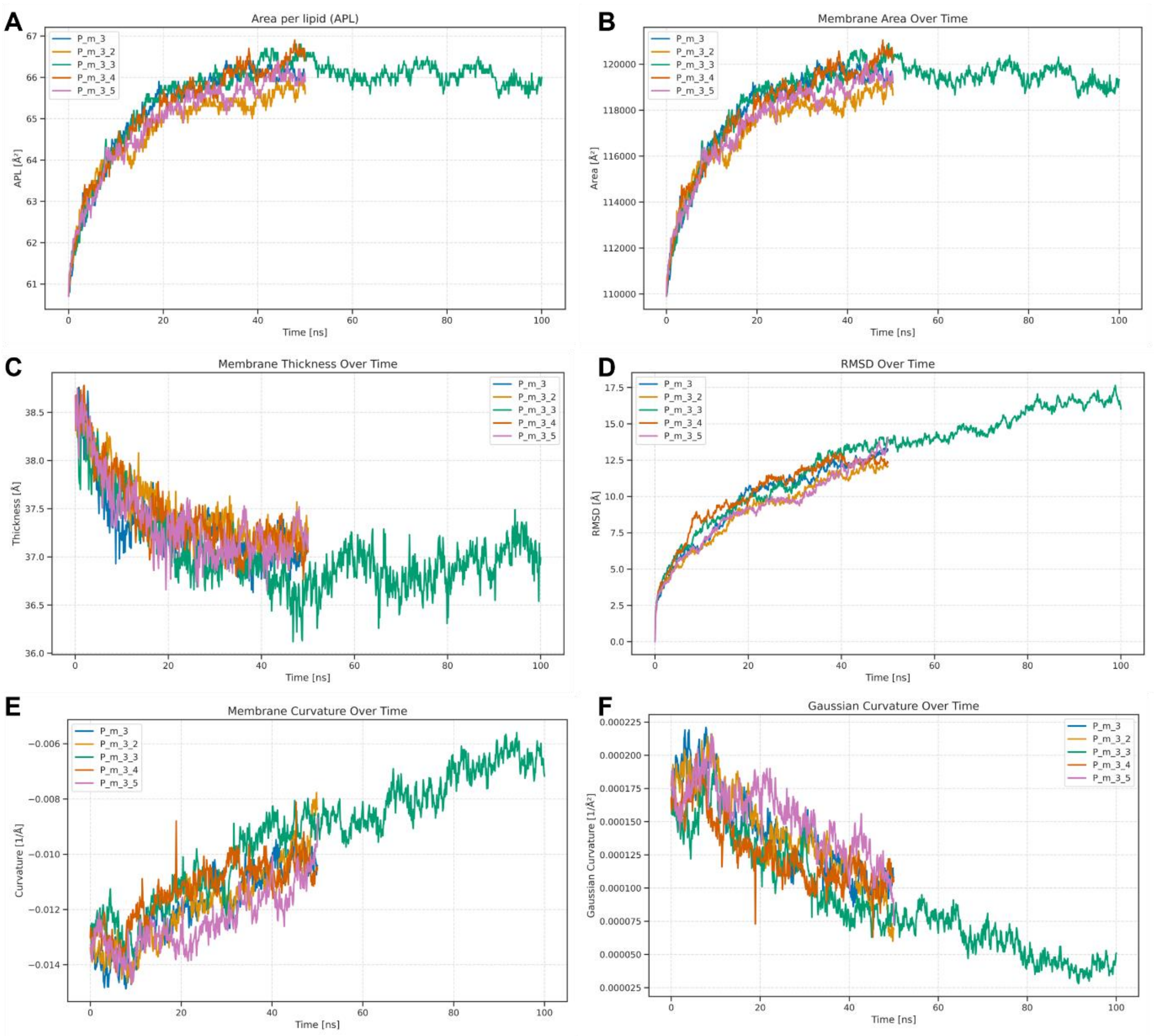


**Fig. S12:** Properties of P_m_3.

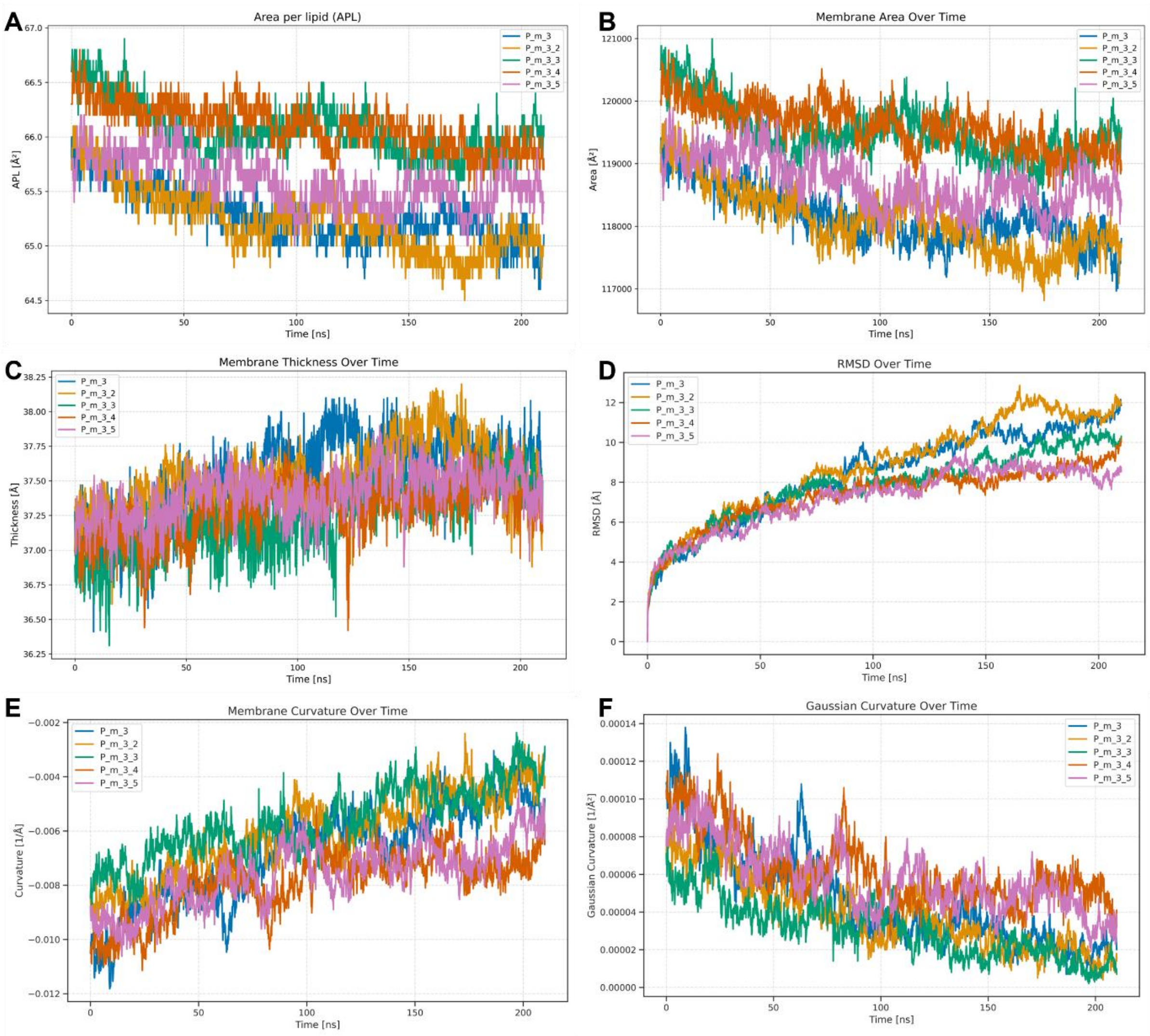


**Fig. S13:** Properties of P_m_3_cef.

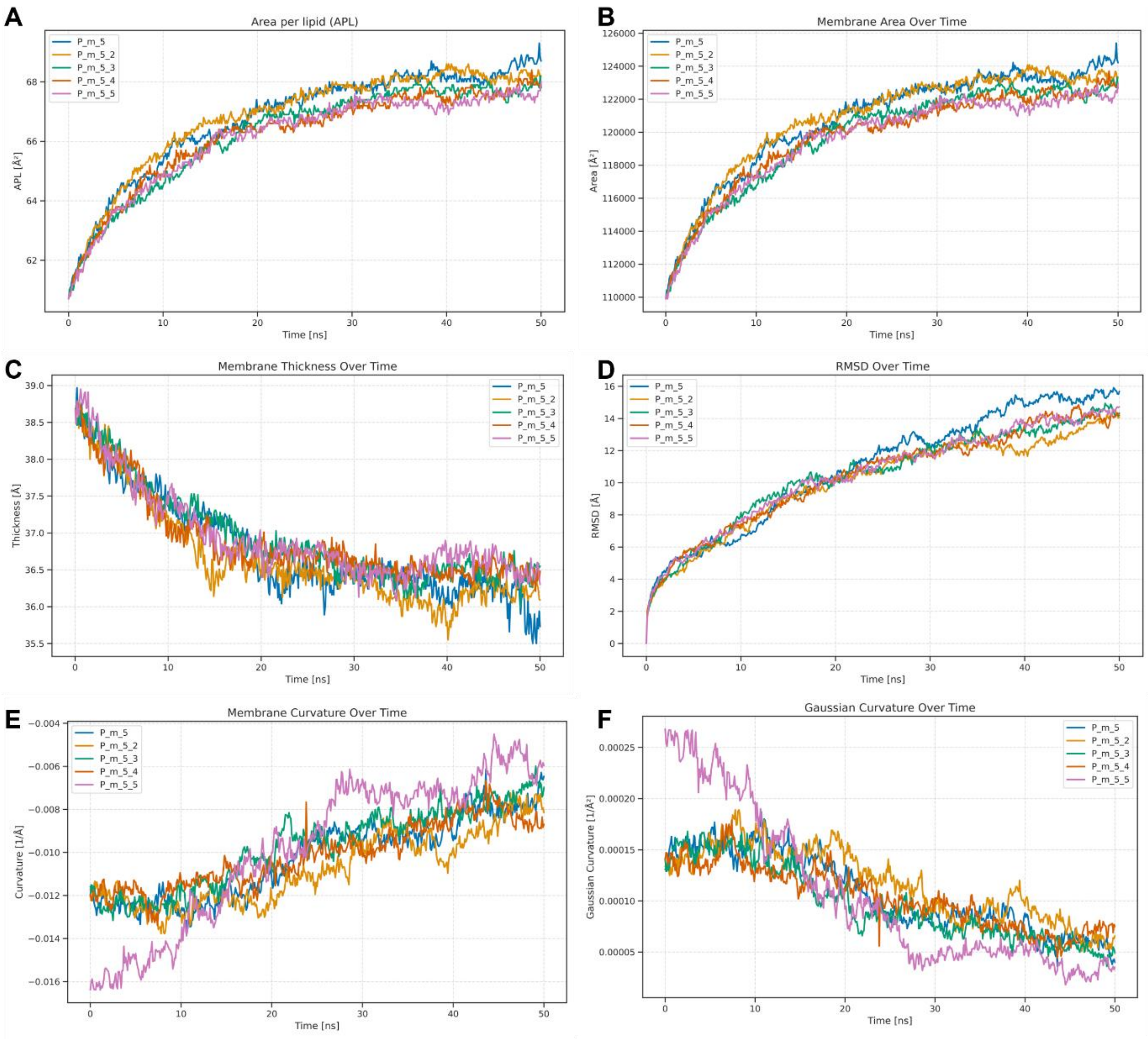


**Fig. S14:** Properties of P_m_5.

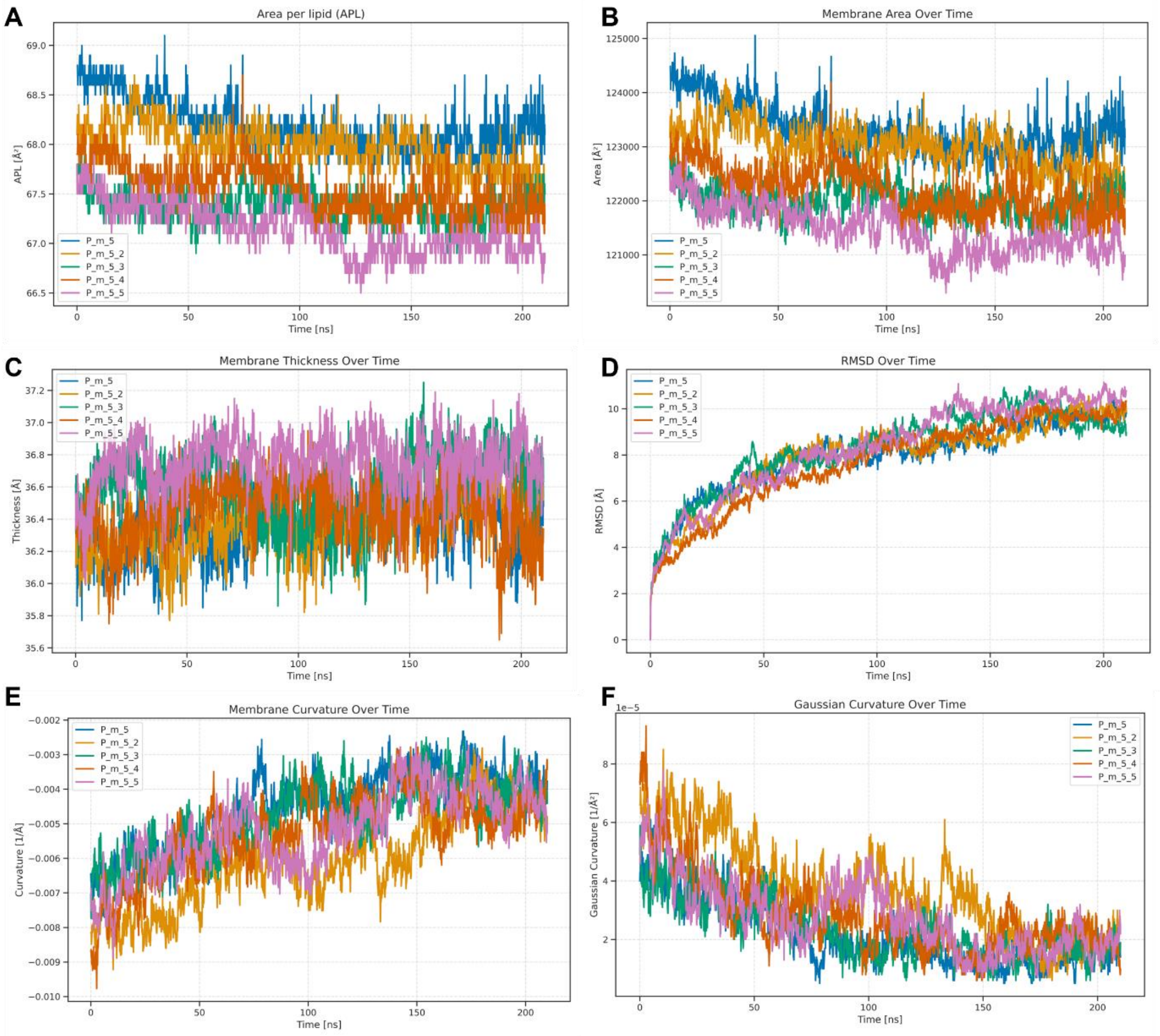


**Fig. S15:** Properties of P_m_5_cef.

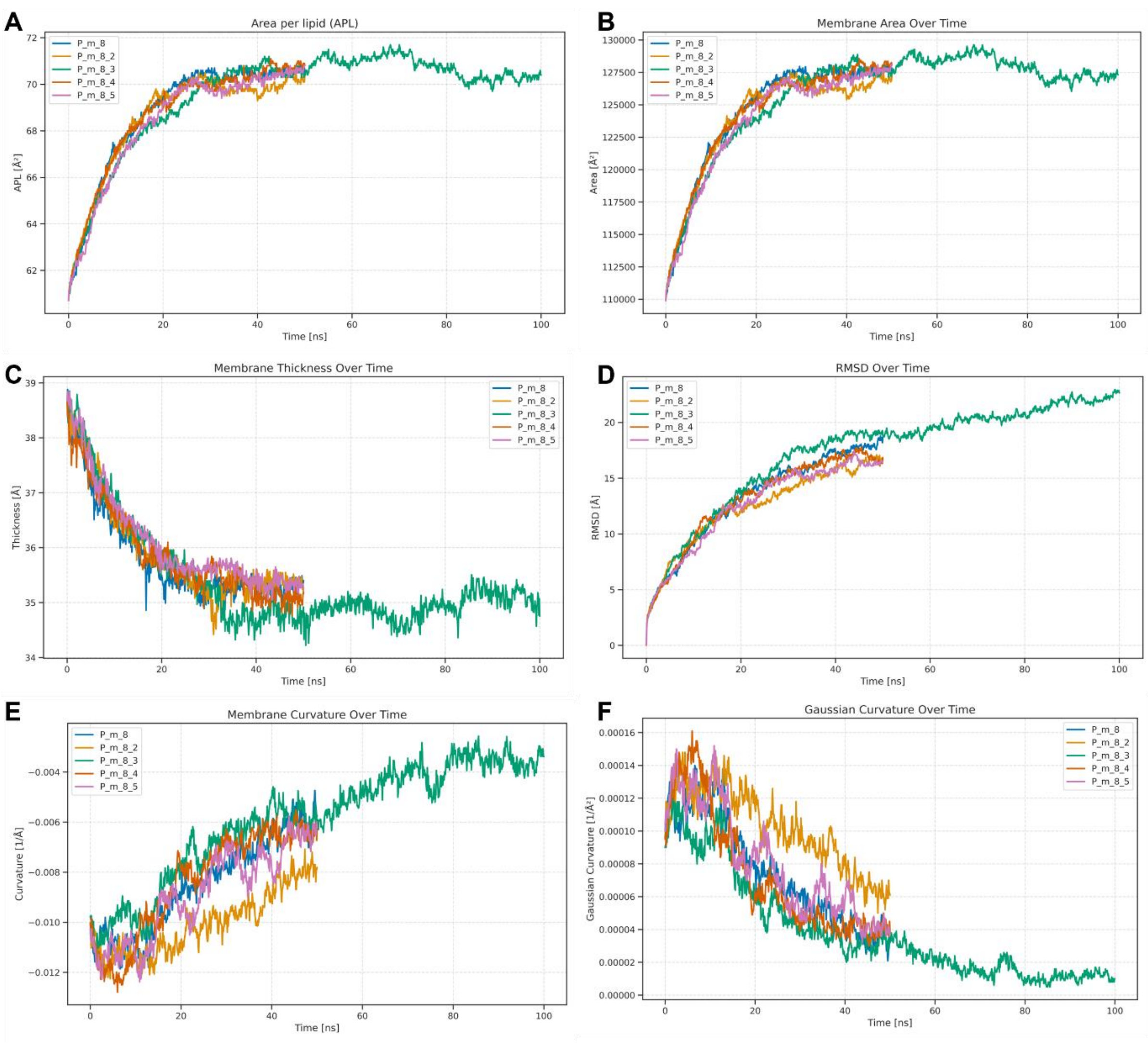


**Fig. S16:** Properties of P_m_8.

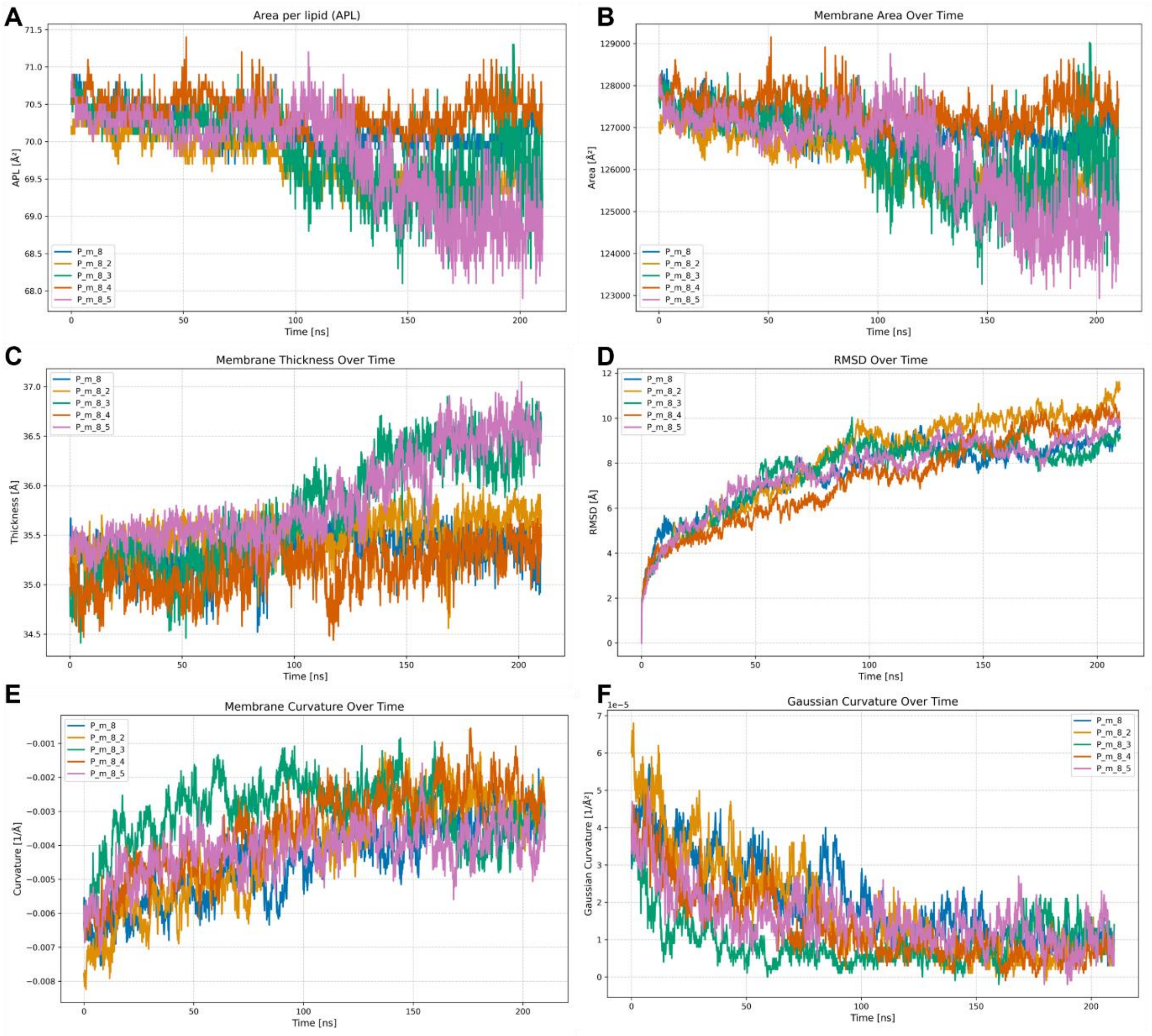


**Fig. S17:** Properties of P_m_8_cef.

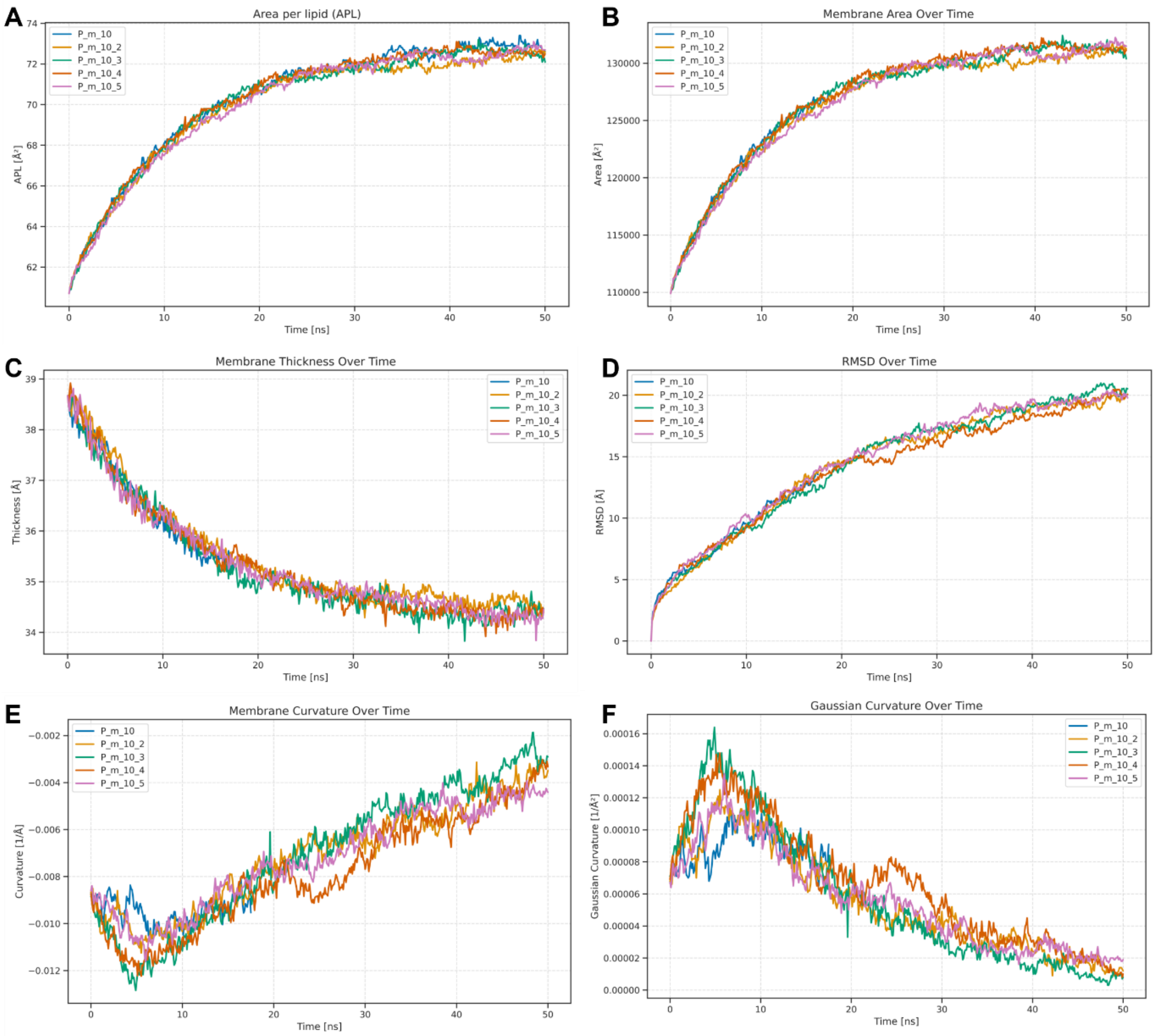


**Fig. S18:** Properties of P_m_10.

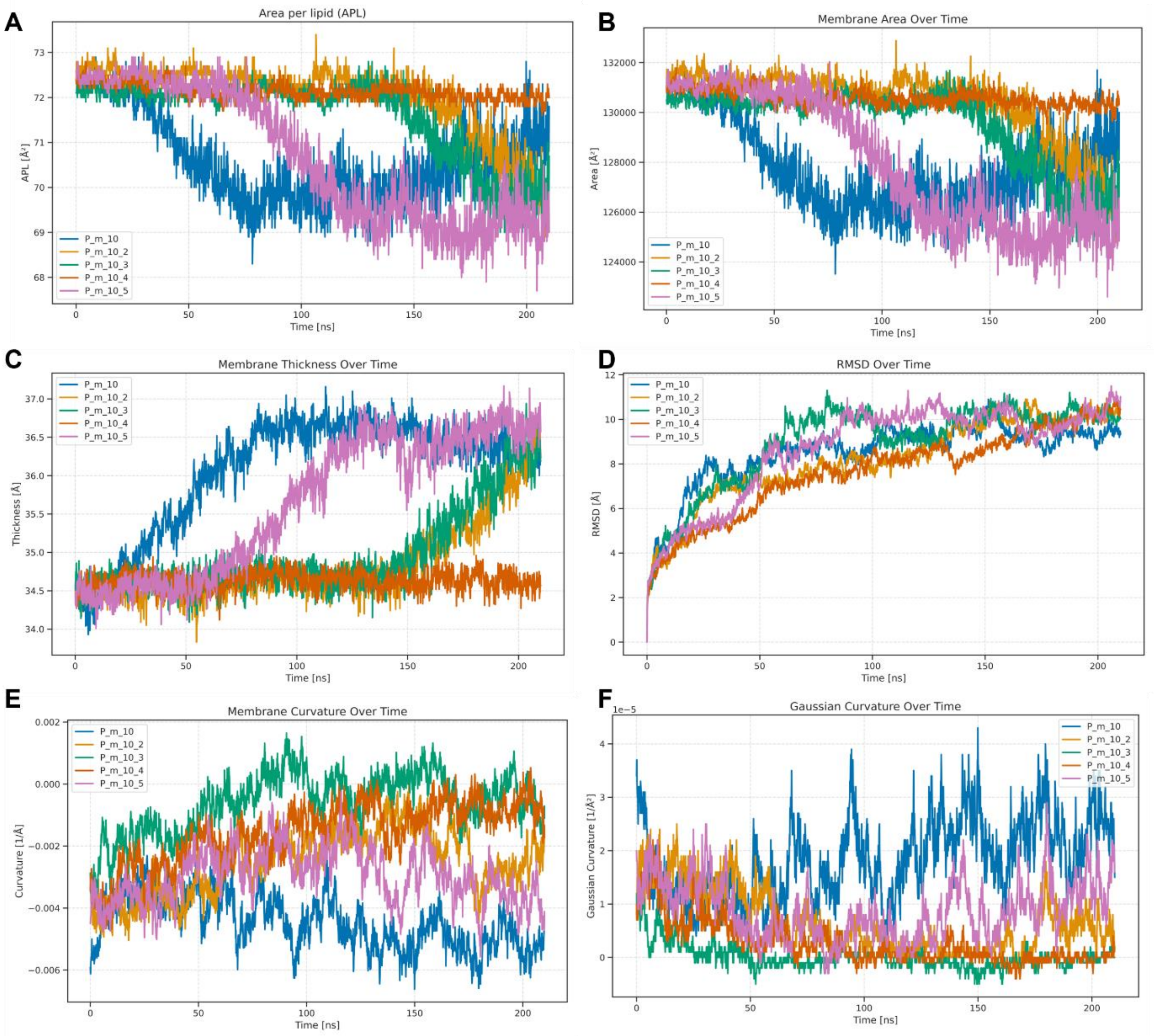


**Fig. S19:** Properties of P_m_10_cef.

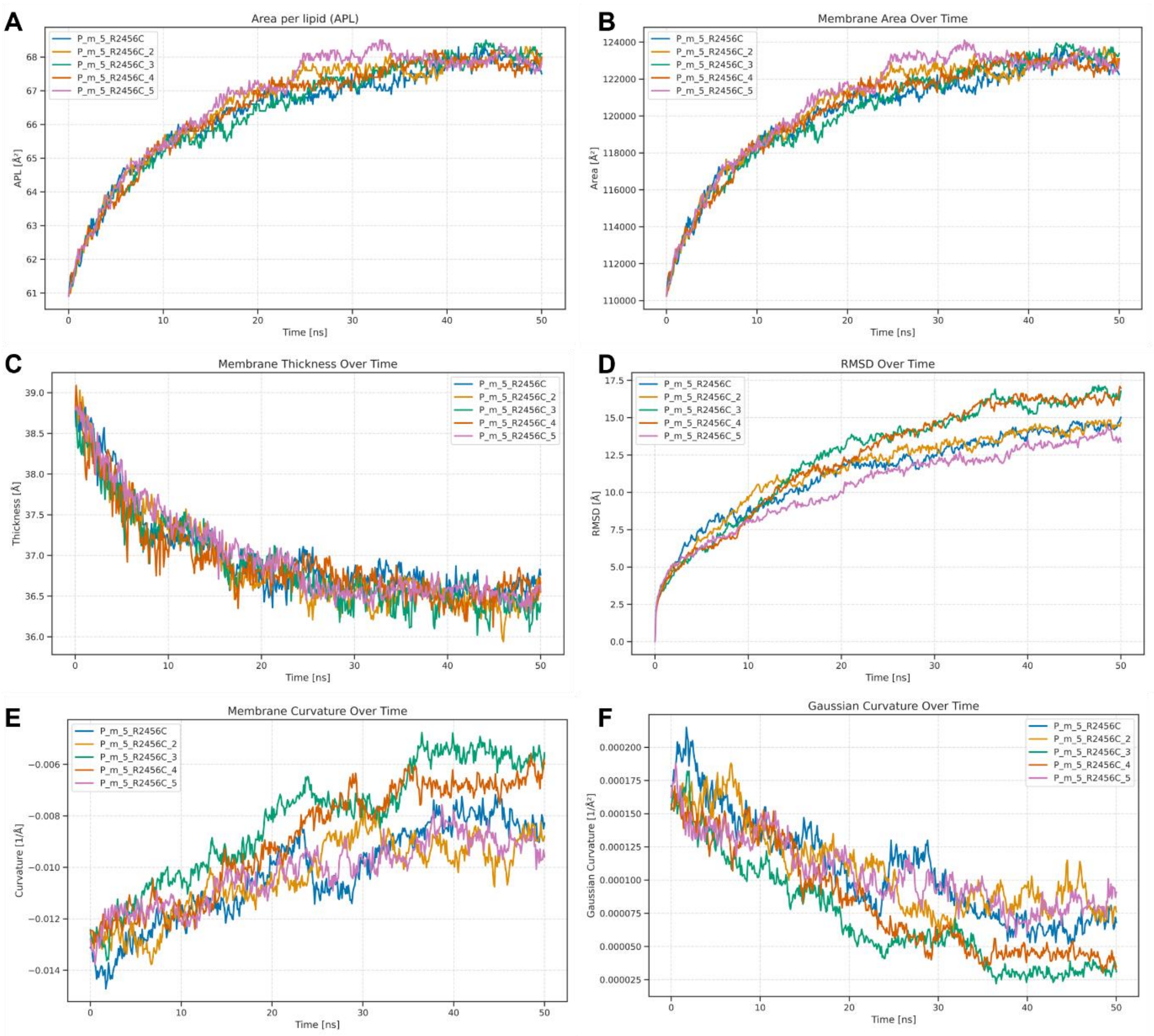


**Fig. S20:** Properties of R2456C.

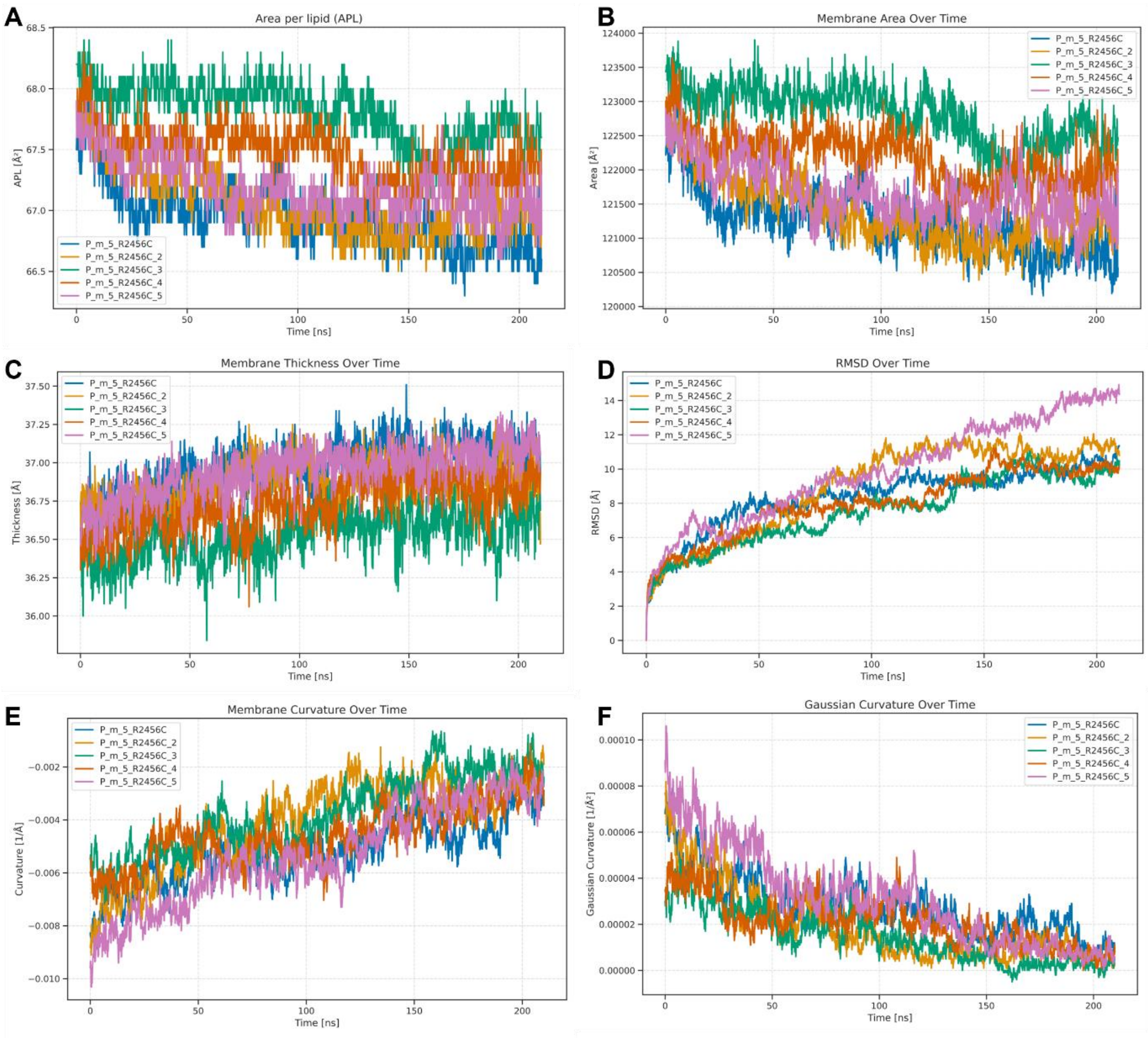


**Fig. S21:** Properties of R2456C_cef.

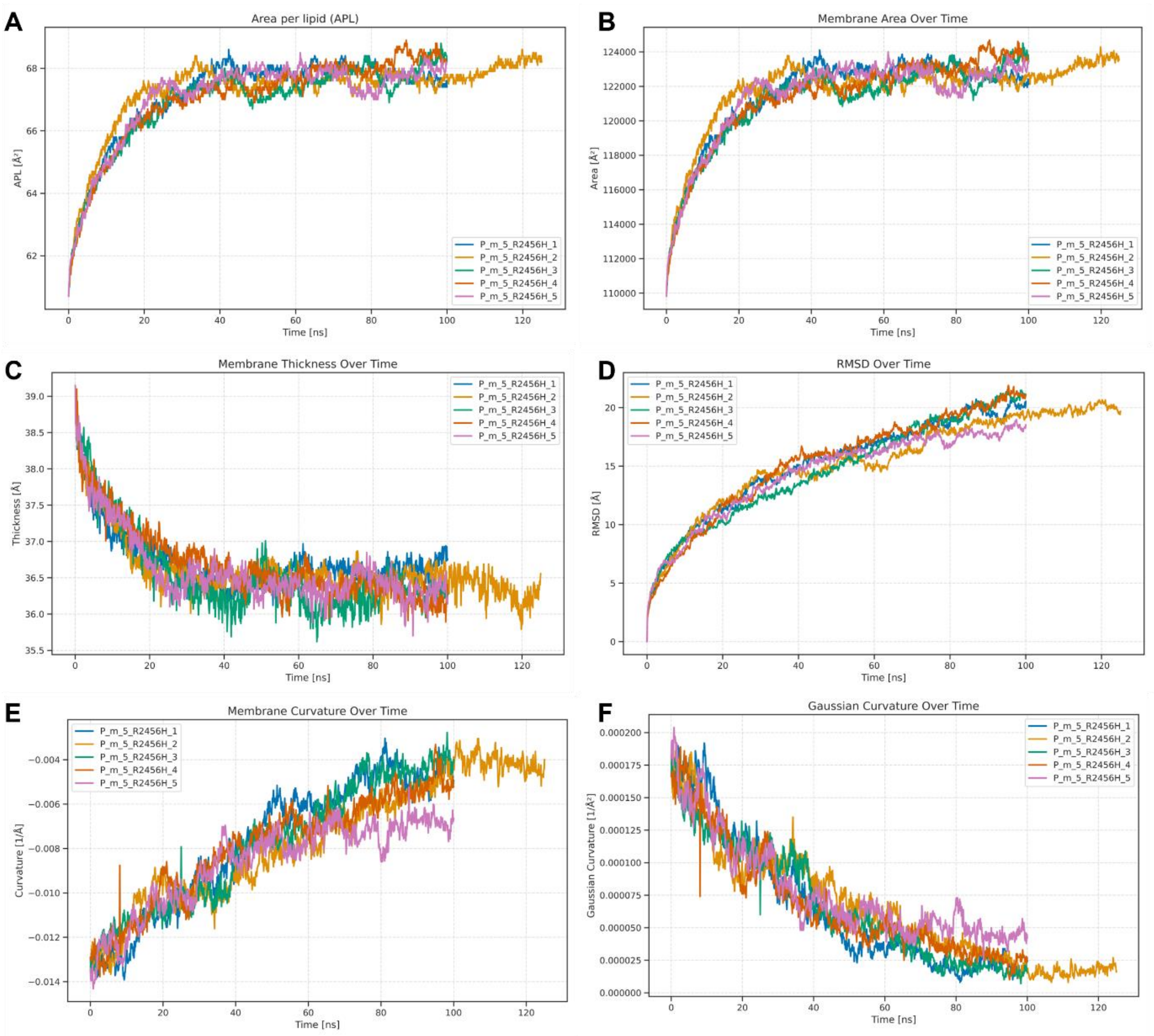


**Fig. S22:** Properties of R2456H.

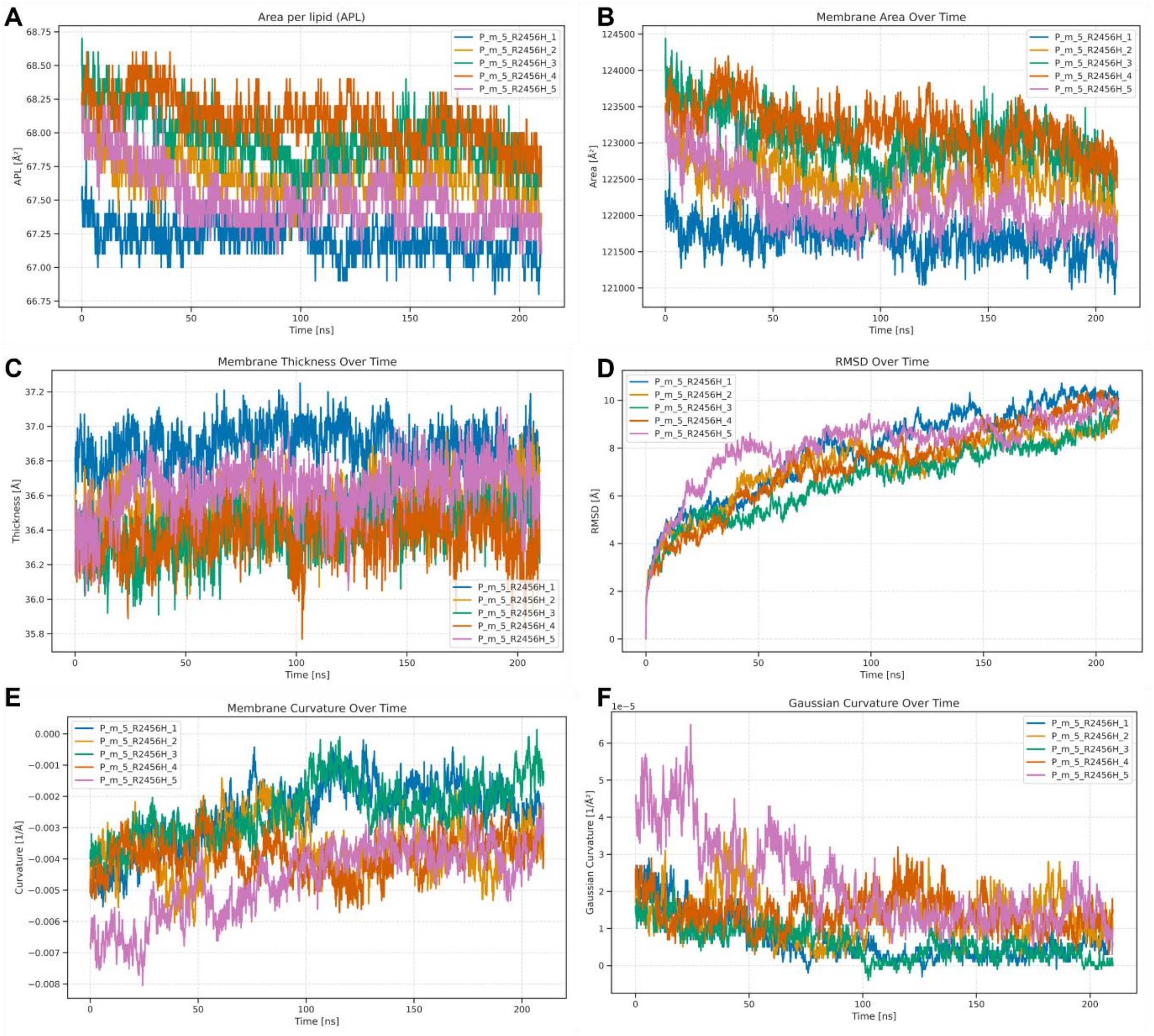


**Fig. S23:** Properties of R2456H_cef.

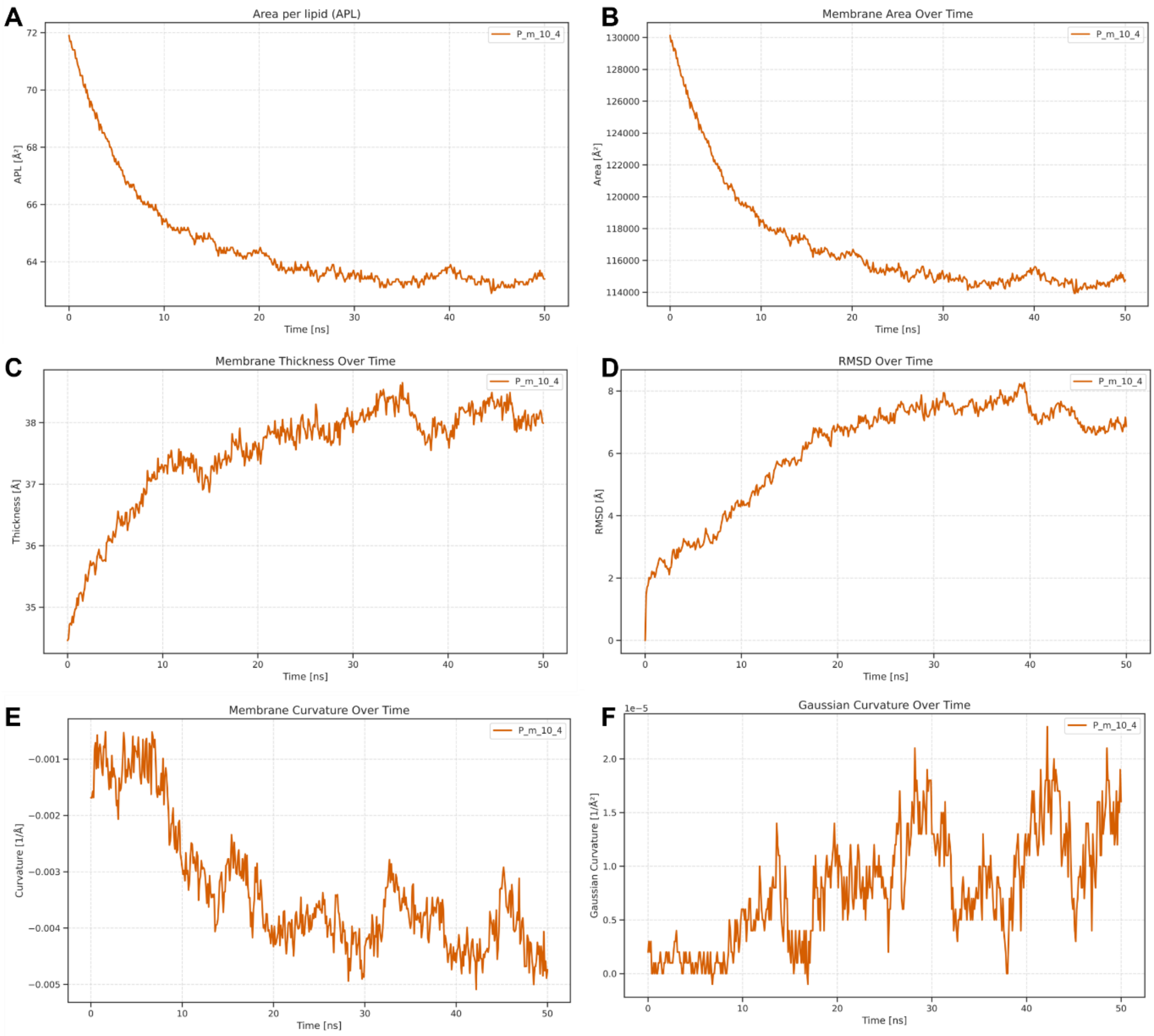


**Fig. S24:** Properties of P_m_10_1bar.

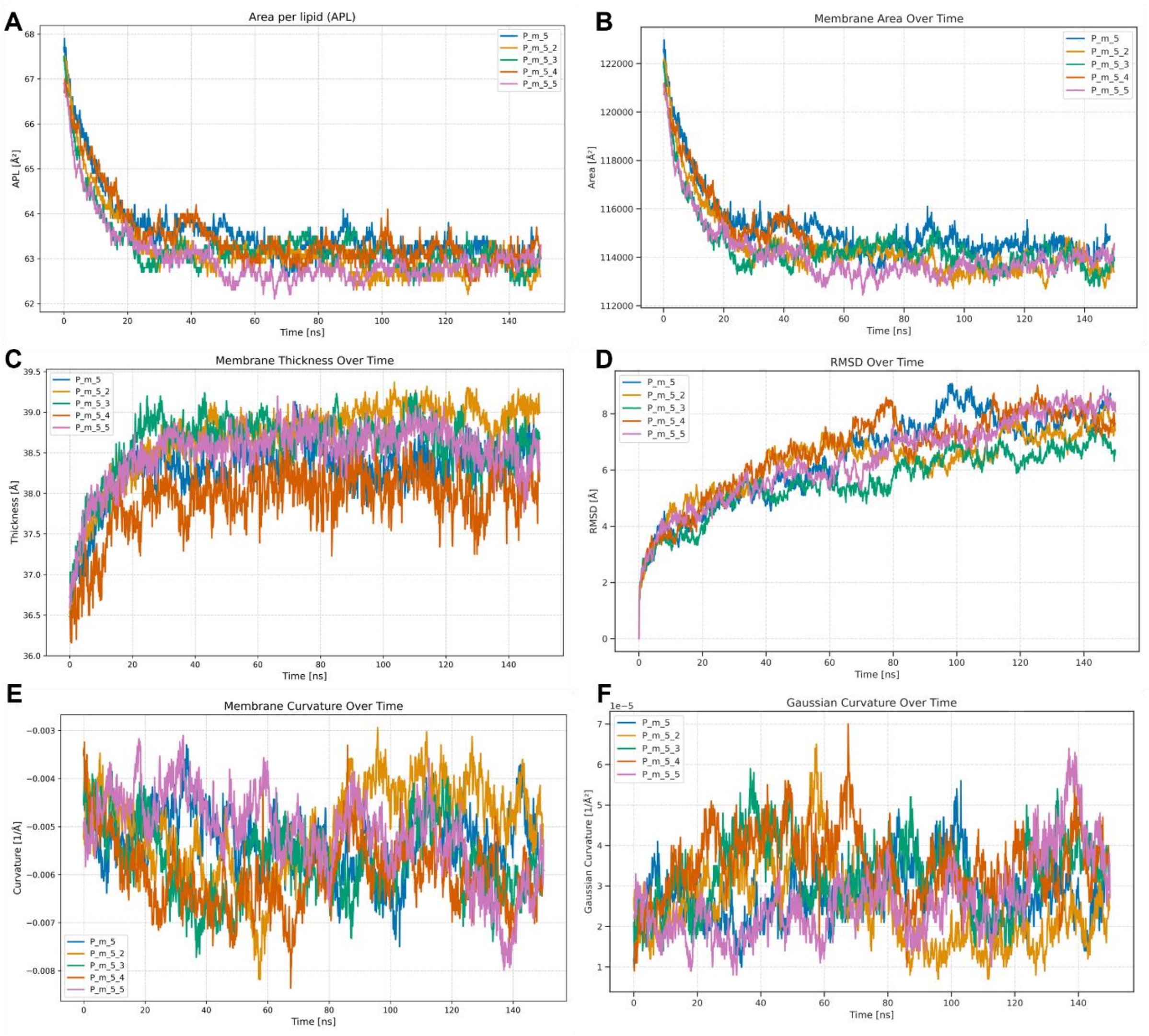


**Fig. S25:** Properties of P_m_5_1bar.

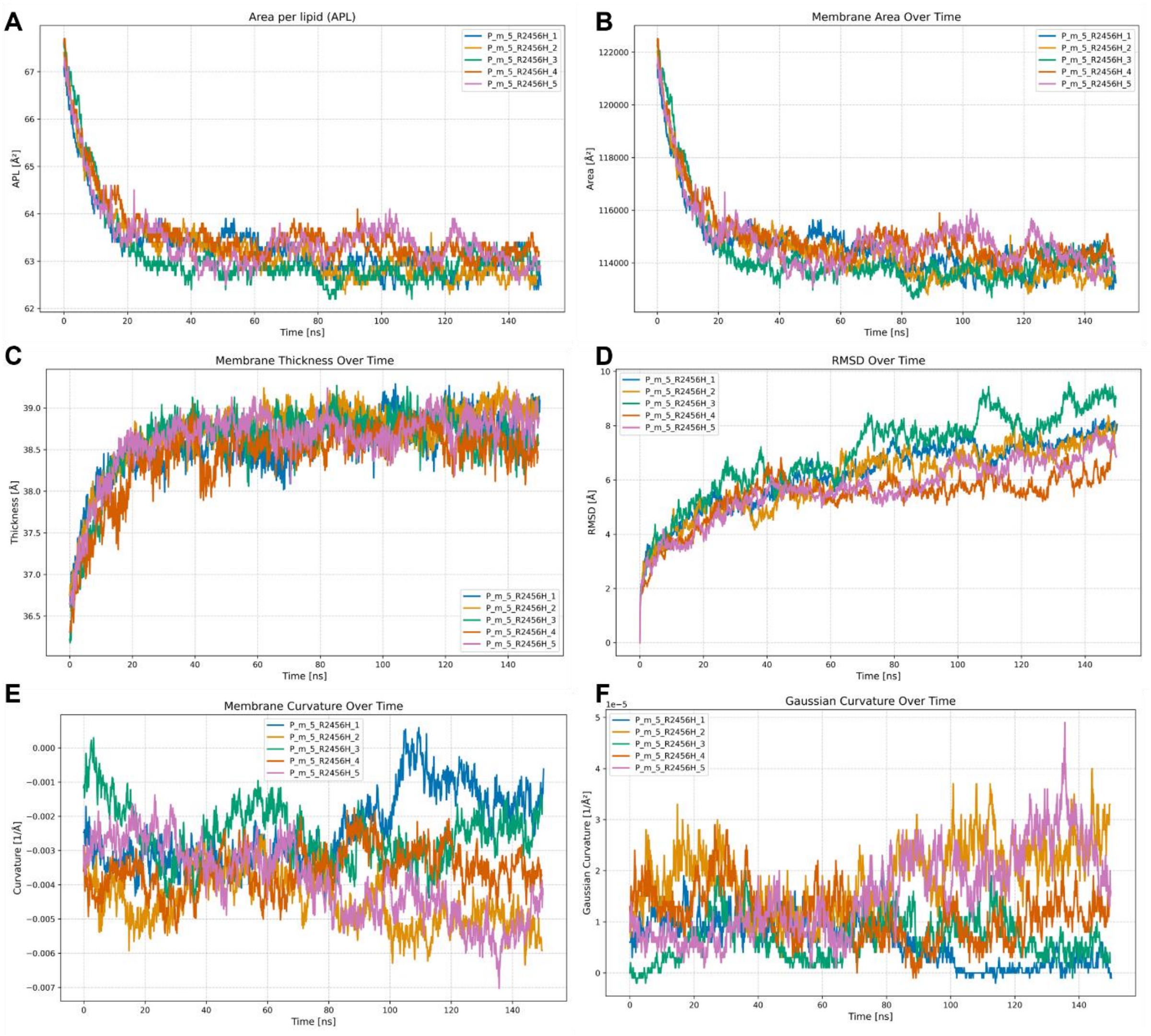


**Fig. S26:** Properties of R2456H_1bar.

**Table S4:** Unsuccessful system setups.

| Protein | Membrane Type | Created with | Force Field | Dims [Å$^3$] (Box Shape) | Atoms | Reason for failure |
|---|---|---|---|---|---|---|
| Mouse PIEZO1 [a] | CHL:POPE:POPC:POPS, (2:2:1:1) Flat | CharmmGUI | Amber | 215x215x187 (Rectangular) | 877,912 | Equilibration failed |
| Human PIEZO1 [b] | CHL:POPE:POPC:POPS, (2:2:1:1) Flat | CharmmGUI | Amber | 280x280x212 (Rectangular) | 1,693,223 | Equilibration failed |
| Mouse PIEZO1 [a] | CHL:POPE:POPC:POPS (2.6:2.4:1.6:1) Curved (r = 250 A) | Packmol-memgen | Amber | 264x264x191 (Rectangular) | 1,494,372 | Equilibration failed |
| Mouse PIEZO1 [a] | POPE Vesicle | Packmol-memgen | Amber | 311x311x311 (Trunctated Octahedron) | 1,590,124 | Equilibration failed |
| Human PIEZO1 [b] | POPE Vesicle | Packmol-memgen | Amber | 348x348x348 (Trunctated Octahedron) | 2,021,310 | Equilibration failed |
| Mouse PIEZO1 [a] | CHL:POPE:POPC:POPS (2.6:2.4:1.6:1) Buckle | Packmol-memgen | Amber | 245x245x294 (Rectangular) | 1,864,405 | Equilibration failed |
| Human PIEZO1 [b] | CHL:POPE:POPC:POPS (2.6:2.4:1.6:1) Buckle | Packmol-memgen | Amber | 295x295x295 (Cubic) | 2,754,683 | Equilibration failed |
| Mouse PIEZO1 [a] | POPE:POPC:POPS (2.4:1.6:1) Buckle | Packmol-memgen | SIRAH | 252x251x314 (Rectangular) | 251,221 | No back-mapping procedure available at the time |
| Human PIEZO1 [b] | POPE:POPC:POPS (2.4:1.6:1) Buckle | Packmol-memgen | SIRAH | 252x251x314 (Rectangular) | 396,873 | No back-mapping procedure available at the time |

[a] based on PDB ID 7WLT[6] [b] Homology model based on PDB ID 6KG7[7] (mouse PIEZO2)
